# Development of a Fluorescent RNA Biosensor for Dual Detection of cGAMP and c-di-GMP Signals in Live Bacteria

**DOI:** 10.64898/2026.01.30.702909

**Authors:** Madeline M. Mumbleau, Ming C. Hammond

## Abstract

Cyclic dinucleotide (CDN) signaling molecules, such as cyclic di-GMP (c-di-GMP) and 3’,3’ cyclic GMP-AMP (cGAMP), are second messengers that play critical roles in phenotypic regulation, such as biofilm formation, host colonization, and bacterial virulence. Recently, hybrid promiscuous (Hypr) GGDEF proteins have been identified in certain bacteria to produce both cyclic dinucleotides. One such enzyme, Bd0367, from the predatory *Bdellovibrio bacteriovorus*, switches between synthesizing c-di-GMP and cGAMP to regulate the bacterial predation cycle and prey exit. However, the molecular mechanism controlling this switch remains unknown. Here, we introduce an RNA-based ratiometric, dual metabolite biosensor that enables simultaneous detection of c-di-GMP and cGAMP in live cells. This sensor integrates a Pepper-based biosensor for c-di-GMP detection and a Spinach2-based biosensor for cGAMP detection into a single transcript, producing distinct fluorescent outputs. In *E. coli*, the dual metabolite sensor reliably reported shifts in c-di-GMP/cGAMP production ratios from various CDN synthases, including Bd0367. Additionally, a histidine kinase was discovered as the probable regulatory partner of Bd0367. These findings demonstrate the sensor’s capacity to assess relative CDN levels and to uncover complex signaling pathways. Together, this ratiometric dual metabolite biosensor provides a foundation for broader applications of fluorogenic RNA biosensors in dissecting bacterial signaling networks, microbial ecology, and host-pathogen interactions.

## INTRODUCTION

Real-time fluorescent monitoring of cellular signaling and other spatiotemporally controlled dynamics has enabled powerful discoveries in the biological sciences (1, 2). Traditionally, molecular sensing and tracking have largely relied on genetically encoded FRET-based biosensors, in which two fluorescent proteins are appended to a target-binding domain to report ligand-induced conformational changes through altered energy transfers between the fluorophores. However, these fluorescent protein-based biosensors are limited in the availability of suitable target-binding domains, are challenging to engineer, and often exhibit low signal-to-noise ratios (3–6).

Fluorogenic RNA aptamers offer an alternative solution for imaging molecular dynamics in live cells, combining minimally invasive tagging with low background fluorescence. Our lab and others have engineered fluorescent RNA-based biosensors with these fluorogenic RNA aptamers that respond to various targets, including cofactors (7–9), metabolites (9–11), signaling molecules (12–16), drugs (9), small RNAs (17–20), and ions (21). Many use naturally occurring, well-structured RNA riboswitches that selectively recognize small molecules and cellular signals, undergoing conformational changes to regulate gene expression (22). When fused to a fluorogenic RNA aptamer, a riboswitch can yield a sensor that fluoresces only upon ligand binding.

The recently developed Pepper RNA aptamer exhibits robust intracellular folding, bright *in vivo* fluorescence, and tunable emission from cyan to red using different synthetic HBC dye analogs without altering the RNA sequence (23). Pepper also does not contain a G-quadruplex motif in its structure, avoiding structural dependence on K^+^ concentrations (24, 25). These properties have led to the development of bright, robust Pepper-based RNA biosensors to detect and measure the abundance of intracellular molecules, such as S-adenosylmethionine (SAM), tetracycline, guanine, and Ag^+^(26, 27), and support multiplexed imaging for simultaneous detection of multiple targets through distinct fluorescent emission profiles.

While protein-based biosensors have been widely developed for single-analyte detection, engineering a genetically encoded dual-analyte protein sensor remains a significant challenge (28). Domain compatibility issues, folding interference, and signal crosstalk have hindered the advancement of dual-analyte protein sensors. As a result, most dual-analyte detection strategies using protein-based biosensors rely on the co-expression of two separate sensors, with each reporting independently (29–31). In contrast, nucleic acid-based sensors offer a greater modality and design flexibility within a single genetic construct, as seen with dual-analyte DNA-based nanodevices that simultaneously sense pH and ion levels in a single assembly with distinct fluorescent outputs (32, 33). To our knowledge, comparable RNA-based sensors with dual functionality within a single transcript have not yet been reported.

Here, we report a novel dual-color, dual metabolite RNA-based biosensor for the bacterial second messengers, cyclic di-GMP (c-di-GMP) and 3’,3’ cyclic GMP-AMP (cGAMP). Before creating the dual metabolite sensor, a previously published second-generation c-di-GMP sensor, Pl-B Spinach2 (13), was adapted to replace the Spinach2 fluorogenic RNA aptamer with the bright, multispectral Pepper aptamer. Optimization was done concurrently *in vitro* and *in vivo,* showing that the Pepper substitution does not disrupt the Pl-B riboswitch function or ligand specificity. This provided the foundation for a ratiometric sensor that contains Pl-B Pepper for c-di-GMP detection and Gm0970 Spinach2 for cGAMP detection, combined within a single transcript, each producing a distinct fluorescence output for simultaneous monitoring of two metabolites in live cells.

The dual biosensor was used to investigate Bd0367, an intriguing hybrid promiscuous (Hypr) GGDEF domain enzyme from the predatory bacterium *Bdellovibrio bacteriovorus* that switches between synthesizing c-di-GMP and cGAMP to regulate distinct aspects of its predatory life cycle (34). The mechanism that controls the enzymatic switch of Bd0367’s product specificity is still unknown. Understanding the regulation Bd0367 is important, as this bacterium preys on and can kill a variety of Gram-negative prey, including pathogens of humans, plants, and animals, generating interest as a potential biocontrol agent and in combating antimicrobial resistance (35–39).

Application of the dual metabolite sensor in *E. coli* allowed for detecting shifts in c-di-GMP/cGAMP production ratios from various CDN synthases, including Bd0367. The sensor was then utilized to screen candidate histidine kinases from *B*. *bacteriovorus* that may interact with Bd0367’s receiver domain and modulate its activity. One histidine kinase candidate was identified that shifts Bd0367 activity to favor c-di-GMP production, and mutational analysis of this kinase suggests a potential regulatory mechanism. These findings demonstrate the biosensor’s capacity to uncover novel enzyme-regulator pathways and highlight its broader application in near real-time dual-signal tracking for studying complex bacterial signaling pathways and the dynamic relationship between two metabolites.

## MATERIAL AND METHODS

### General Reagents and Oligonucleotides

All DNA oligonucleotides used for constructs and cloning were purchased from the University of Utah HSC Core facility. DFHBI-1T was purchased from Sigma-Aldrich (St. Louis, MO), and HBC dyes were purchased from either FR Biotechnology (Shanghai, China) or MedChemExpress (Monmouth Junction, NJ) and were both stored in DMSO as a 50 mM stock for DFHBI-1T and a 100 µM stock for HBC dyes. Chemically competent BL21 (DE3) Star cells were purchased from the Berkeley MacroLab (Berkeley, CA) for all *in vivo* Pl-B biosensor, Dual Biosensor, and histidine kinase analyses.

### *In Vitro* Transcription

For *in vitro* biosensor screening experiments, DNA templates for *in vitro* transcription were prepared by PCR amplification using Phusion DNA polymerase (Berkeley MacroLab) from pET31b plasmids containing the constructs. The forward primer includes an extended T7 promoter, and the reverse primer is specific to the 3’ end of the Pepper biosensor constructs. After the PCR reaction, products were treated with Dpn1 at 37 °C for 1 h to digest the plasmid template before purification by QIAquick PCR purification kit (Qiagen).

RNA was transcribed from DNA templates following a method similar to that previously described (40). Briefly, ∼1-1.5 µg of DNA template is incubated for 4 h at 37 °C with homemade T7 RNA polymerase in 40 mM Tris-HCl, pH 8.0, 6 mM MgCl_2_, 2 mM spermidine, 10 mM DTT, 1 U of inorganic pyrophosphate, and 2 mM rNTPs. RNAs are purified by separation on a 6 % denaturing PAGE gel (7.5 M urea), visualized by UV shadowing, and extracted from gel pieces using Crush Soak buffer (10 mM Tris-HCl, pH 7.5, 200 mM NaCl, and 1 mM EDTA, pH 8.0). Purified RNAs are precipitated from Crush Soak buffer with ethanol, dried, and resuspended in ddH_2_O. Accurate RNA concentrations are determined by performing the hydrolysis assay to eliminate the hypochromic effect due to RNA secondary structure (41).

### General Procedure for *In Vitro* Fluorescence Assays

*In vitro* fluorescence assays were carried out in a binding buffer containing 40 mM HEPES (pH 7.5), 125 mM KCl, and 3 mM MgCl_2_ at 28 °C or 37 °C. Other conditions, including temperature and the concentrations of DFHBI-1T and HBC dyes (HBC485, HBC514, or HBC620) and RNA, varied as indicated in the main text and supplementary information figures. The RNA was renatured in 2x binding buffer at 72 °C for 3 min and slowly cooled to ambient temperature for 10 min before being added to the binding reaction. HBC dyes (or water for no dye control) were added to the solution containing binding buffer first, followed by RNA, and incubated for 1 h before fluorescent measurement.

Fluorescent measurements were performed in 30 µL or 300 µL volumes, and fluorescence emission was recorded in a Greiner Bio-One 384-well black plate using a SpectraMax i3x plate reader (Molecular Devices) for 10 min. The fluorescence emission was calculated as an average (n=3) of endpoint values using the following parameters: (HBC485) 443 nm excitation, 485 nm emission; (DFHBI-1T) 480 nm excitation, 505 nm emission; (HBC514) 458 nm excitation, 514 nm emission; (HBC620) 577 nm excitation, 620 nm emission.

For Pl-B Pepper biosensor ligand specificity analysis, 100 nM of Pl-B Pepper biosensor RNA was incubated with 200 nM of HBC514 with 25 µM of various ligands. Fluorescence was normalized to Pl-B Pepper with 3’3’ c-di-GMP. Fluorescence turn-on was calculated by dividing the fluorescence in the presence of cyclic di-GMP by the fluorescence in the absence of cyclic di-GMP.

For dual biosensor ligand specificity analysis, 100 nM of Dual Biosensor RNAs were incubated with 10 µM of DFHBI-1T, 200 nM of HBC620, and 1 µM of cyclic dinucleotides, c-di-GMP and cGAMP. Fluorescence activation for each individual sensor was calculated by dividing the fluorescence in the presence of cyclic dinucleotides by fluorescence in the absence of cyclic dinucleotides.

### Binding Affinity Analysis of Cyclic di-GMP and Cyclic GMP-AMP Biosensors

The binding affinities of Pl-B Pepper biosensors were measured with 20 nM RNA in a binding buffer containing 100 nM HBC514, 40 mM HEPES (pH 7.5), 125 mM KCl, and 3 mM MgCl_2_ at 28 °C or 37 °C. The cyclic di-GMP concentration was varied from 10 pM to 25 µM. The fluorescence of the binding buffer was subtracted as background to determine relative fluorescence units. The dissociation constant (*K*_d_) for each binding event was calculated from the concentration-dependent fluorescence curves by fitting the normalized fluorescence intensity (*F*_N_) versus the log of cyclic-di-GMP concentration plot to a nonlinear regression (log (agonist) versus response (three-parameter)) using Prism 10 software. *F*_N_ was calculated as (*F_i_* – *F*_0_)/(*F*_s_ – *F*_0_), where *F_i_* is fluorescence intensity at each ligand concentration, *F*_0_ is fluorescence intensity without ligand, and *F*_s_ is fluorescence intensity at the saturation point.

The binding affinities of optimized Pl-B Pepper and Gm0970 Spinach2 biosensors were measured with 20 nM RNA in a binding buffer containing 10 µM DFHBI-1T or 200 nM HBC514, 40 mM HEPES (pH 7.5), 125 mM KCl, and 3 mM MgCl_2_ at 28 °C or 37 °C. The cyclic di-GMP and 3’3’-cGAMP concentrations were varied from 1 nM to 50 µM. The fluorescence of the binding buffer was subtracted as background to determine relative fluorescence units. The dissociation constant (*K*_d_) for each binding event was calculated from the concentration-dependent fluorescence curves by fitting the normalized fluorescence intensity (*F*_N_) versus the log of the specified cyclic di-nucleotide concentration plot to a nonlinear regression (log (agonist) versus response (three-parameter)) using Prism 10 software. *F*_N_ was calculated as (*F_i_* – *F*_0_)/(*F*_s_ – *F*_0_), where *F_i_* is fluorescence intensity at each ligand concentration, *F*_0_ is fluorescence intensity without ligand, and *F*_s_ is fluorescence intensity at the saturation point.

### Biosensor and Synthase Cloning

Pl-B Pepper biosensors were appended with a tRNA scaffold constructs, for *in vivo* expression, through overhang addition by PCR, the resulting products were subcloned into the pET31b vectors via a two-piece Gibson, and were assembled with the Gibson Assembly Master Mix (New England Biolabs, Ipswich, MA, USA), where both linear backbone and insert fragments were amplified with either Phusion or Q5 DNA polymerase (New England Biolabs). WspR, cyclic diguanylate synthase enzymes, was cloned into pCOLADUET-1 vectors through a two-piece Gibson Assembly cloning, for which both linear backbone and insert fragments were amplified with Phusion.

Dual biosensors were appended with the T7 promoter and T7 terminatory through overhang addition by PCR, the resulting products were subcloned into the pET31b vectors via a two-piece Gibson were assembled with the Gibson Assembly Master Mix (New England Biolabs, Ipswich, MA, USA), where both linear backbone and insert fragments were amplified with either Phusion or Q5 DNA polymerase (New England Biolabs). Cyclic dinucleotide synthase enzymes and histidine kinases were cloned into pCOLADUET-1 vectors through a two-piece Gibson Assembly cloning, for which both linear backbone and insert fragments were amplified with Phusion. Histidine kinase candidates were codon-optimized for *E. coli* and cloned into pET28a(+) plasmids by Twist Biosciences (San Francisco, CA, USA). Insertion of cyclic dinucleotide synthases and histidine kinases into the desired plasmid was achieved with primer overhangs complementary to the plasmid backbone. pCOLADUET-1 vectors containing both cyclic dinucleotide synthases and histidine kinase candidates are cloned in divergent directions. Cloning Site 1: Histidine Kinases (Reverse), Cloning Site 2: Cyclic Dinucleotide Synthases (Forward).

### *In Vivo* Flow Cytometry

Chemically competent *E. coli* BL21 (DE3) Star cells were co-transformed with plasmids containing Pl-B Pepper constructs flanked by a tRNA scaffold and WspR synthases using heat shock, and cells were plated on LB/carbenicillin/kanamycin plates (Carb: 50 µg/mL; Kan: 50 µg/mL) for Pl-B Pepper biosensor analyses. Similarly, chemically competent *E. coli* BL21 (DE3) Star cells were co-transformed with plasmids containing Dual Biosensor constructs and Cyclic dinucleotide synthases using heat shock, and cells were plated on LB/carbenicillin/kanamycin plates (Carb: 50 µg/mL; Kan: 50 µg/mL) for Dual Biosensor and histidine kinase discovery studies.

For flow cytometry assays, single *E. coli* colonies (sample size indicated in relevant figures) containing Pepper biosensors or Dual Biosensor and cyclic dinucleotide synthases or histidine kinase- Bd0367 synthase were inoculated in a 2 mL culture of noninducing media (NI) containing carbenicillin (50 µg/mL) and kanamycin (50 µg/mL) for biological replicates. Cultures were grown at 37 °C in an incubator shaking at 250 rpm for 22-24 h. After growth in NI media, the culture was diluted 100x into 3 mL of ZYP-5052 autoinduction media (AI) containing carbenicillin (50 µg/mL) and kanamycin (50 µg/mL) and grown for 16-18 h at 37 °C, shaking at 250 rpm to express biosensor constructs and cyclic dinucleotide synthases and histidine kinases proteins. Cells were then diluted 200x into 1x PBS with 200 nM HBC514 or with 50 µM DFHBI-1T and 200 nM HBC620 and incubated at room temperature for 1 h to ensure full dye equilibration before measuring on the flow cytometer.

For testing Bd0367 synthase dependence on intracellular ATP levels, 5 µM of ATPase inhibitor, carbonyl cyanide m-chlorophenyl hydrazone (CCCP) was added into cells expressing Bd0367 only after 1 h of HBC620 and DFHBI-1T dye incubation in 1x PBS. Cells, dyes, and CCCP were incubated together at room temperature for 30 min before measuring cyclic dinucleotide levels on the flow cytometer. ATPase inhibitor concentration and incubation time were based on the results shown by Diez-Gonzalez et al (42).

For cyclic dinucleotide synthase and histidine kinase studies, single-cell fluorescence was measured on the Beckman Coulter Cytoflex S (Utah Flow Core facility) using the following settings: excitation lasers, 488 nm and 561 nm; emission filters, FITC and PE-TxRed; cell counts for each measurement, 30,000 or 50,000. The ATPase inhibitor and some histidine kinase mutational analyses, single-cell fluorescence were measured using the Attune NxT flow cytometer with an autosampler (Life Technologies) using the following settings: excitation lasers, 488 nm and 561 nm; emission filters, GFP and mCherry; cell counts for each measurement, 50,000. The raw data were analyzed using FlowJo software. Statistical analyses (two tailed *t*-test and identifying outliers via ROUT method; Q= 5%) were performed using Prism 10 software.

Fluorescence activation was calculated by dividing the fluorescence in the presence of cyclic di-GMP or cGAMP, depending on the cyclic dinucleotide synthase expressed, by the fluorescence in the absence of cyclic di-GMP or cGAMP with a catalytically inactive WspR G249A. Dual Biosensor cyclic di-GMP levels were determined by a median fluorescence intensity ratio of MFI_TxRed_/ MFI_FITC_ for each biological replicate. cGAMP levels were determined by a median fluorescence intensity ratio of MFI_FITC_/ MFI_TxRed_ for each biological replicate. Statistical analyses (two tailed *t*-test and identifying outliers via the ROUT method; Q= 5%) were performed using Prism 10 software.

## RESULTS

### *In Vitro* and *In Vivo* Characterization of the Pepper-Based Cyclic di-GMP Sensor

Similar to other RNA-based biosensors, a Pepper-based biosensor is composed of three domains: Pepper as the fluorescent indicator, an RNA aptamer such as a natural riboswitch that binds to ligands with high specificity, and a transducer domain that transmits a fluorescent signal upon a ligand-binding event (12) (Supplementary Figure S1). Initial constructs of the Pl-B Pepper biosensor utilize an inverted version of the Pepper aptamer, known as iPepper (43), fused to the Pl-B riboswitch (13, 44) with four transducer stems (TS1-TS4) (Supplementary Figure S2, Supplementary Table S1) (45). *In vitro* testing showed that constructs with the shorter **TS1** and **TS2** stems exhibit fluorescence increases of 1.3-fold and 3.3-fold, respectively, with 25 µM c-di-GMP, whereas the longer stems were unresponsive. In *E. coli* expressing Pl-B Pepper biosensors flanked by a tRNA scaffold (46), constructs **TS1** and **TS2** exhibited higher fluorescence turn-on (2.4-fold and 1.7-fold) than **TS3** and **TS4** when co-expressed with an active c-di-GMP synthase WspR compared to a catalytically inactive control, WspR G249A (Supplementary Figure S2B). Based on these results, Pl-B Pepper **TS1** and **TS2** were selected for further optimization.

To improve performance, two additional transducer stems, similar in length to **TS1** and **TS2,** were generated (Figure 1). *In vitro* analysis of these new transducers revealed that **TS1.1** showed improved fluorescence turn-on (4.5-fold) with 25 µM c-di-GMP, but **TS2.1** did not (1.6-fold) (Figure 1B). This trend held with blue and red HBC dyes, HBC485 and HBC620 (Supplementary Figure S3). However, in cells co-expressing the Pl-B Pepper constructs with WspR with 200 nM HBC514, both **TS1.1** and **TS2.1** exhibited much more robust fluorescence activation (37.5-fold and 12-fold, respectively) than **TS1** and **TS2** (Figure 1C). Based on high fluorescence responsiveness *in vitro* and in cells, the **TS1.1** construct was chosen for all subsequent experiments and called Pl-B Pepper.

**Figure 1.**
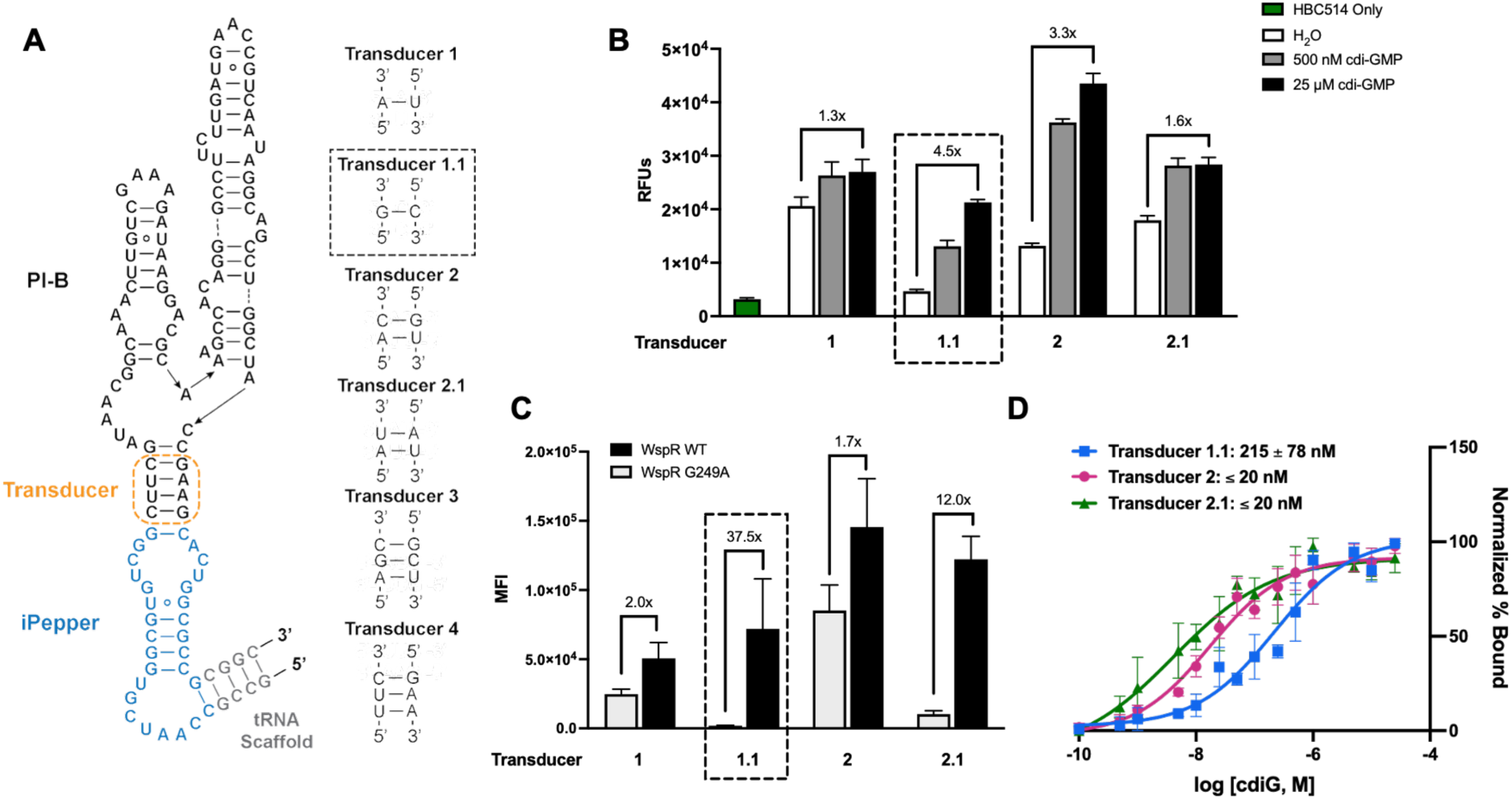
Pl-B Pepper Sensor Optimization and Screening *In Vitro* and *In Vivo*. (A) Sequence and secondary structure model of Pl-B Pepper construct, which comprises the Pl-B riboswitch (black), a transducer domain region (boxed in orange), and the iPepper aptamer (blue). For in vivo studies, Pl-B Pepper sensors are within a tRNA scaffold (gray). A series of transducer domains was tested to optimize fluorescence response by decreasing the fluorescence background signal. (B) Fluorescence activation of Pl-B Pepper sensors and transducer domains in vitro with 0, 500 nM, and 25 µM c-di-GMP and measured as relative fluorescence units (RFUs). HBC514 dye background fluorescence is represented as a green bar. Fluorescence activation levels are based on the fold difference of fluorescence levels with 0 µM c-di-GMP versus 25 µM c-di-GMP. The optimal sensor is indicated with a black dotted box. The data shown are the average with a standard deviation of three independent replicates. (C) Average median fluorescence (MFI) measured by flow cytometry of E. coli BL21 (DE3) Star cells co-expressing biosensor constructs with different transducer domains as indicated above, with various diguanylate cyclase enzymes. Black represents an inactive mutant of diguanylate cyclase WspR (G249A); white represents wild-type WspR. Fluorescence activation is determined with the mean fluorescence of each Pl-B Pepper construct under different enzyme conditions. The optimal sensor is indicated by a black dotted box. Data are from three biological replicates by analyzing 30,000 cells per sample after 1 h incubation with HBC514 dye (200 nM). Data are the average with a standard deviation of three biological replicates. (D) Biosensor binding affinity measurements are shown as normalized % bound (maximum fluorescence), normalized for each biosensor, with titration of c-di-GMP at 3 mM Mg2+, 37 °C. Points and error bars represent averages and standard deviations, respectively, from three independent replicates. Best-fit curves are also shown, and the KDs of each biosensor are listed in the legend.

Pl-B Pepper biosensor was further characterized for its *in vitro* binding affinity and ligand selectivity. Pl-B Pepper displayed a *K*_d_ of 215 ± 78 nM and a maintained responsive range from 10 % to 90 % signal from 1 nM to 1 µM c-di-GMP, which covers well the dynamic range of c-di-GMP levels within bacteria (Figures 1D, Supplementary Figure S4) (45–49). Pl-B sensors also retain selectivity toward 3’3’ c-di-GMP and its synthetic analog, 2’3’ c-di-GMP, over other cyclic di-nucleotides and related compounds (Supplementary Figure S6). Similar to Pl-B Spinach2, Pl-B Pepper sensors respond to high concentrations of pGpG, but the Pl-B Pepper designs are about 120-fold more selective towards c-di-GMP. Furthermore, mutational analysis was performed to demonstrate that the fluorescence response in cells is due to c-di-GMP binding to the Pl-B riboswitch (Supplementary Figure S5). Pl-B Pepper and its mutants were co-expressed in cells with WspR variants: inactive WspR G249A, wild-type WspR, and a constitutively active WspR D70E. Disruption of the P1 stem of the Pl-B riboswitch (M1 mutant) led to a reduced fluorescence response in cells expressing WspR D70E. In contrast, mutants M2 and M3 directly affect the hydrogen bonding to c-di-GMP in the riboswitch binding pocket, and both exhibited minimal to no fluorescence response. Taken together, these results show that the fluorescence of the Pl-B Pepper sensor observed *in vitro* and in cells is selectively activated in response to c-di-GMP.

### Dual Metabolite Sensor Design and *In Vitro* Characterization

Previous work has demonstrated the functionality of two orthogonal fluorogenic RNA aptamer pairs, HBC620-Pepper and DFHBI-1T-Spinach2, within the tRNA-DF30 scaffold (47). This improved scaffold greatly enhanced the fluorescence output of both aptamers and provided a basis for displaying the orthogonal RNA aptamers within a single RNA transcript. Expanding on this work, we set out to develop a dual metabolite biosensor displayed within the tRNA-DF30 scaffold to investigate the function of the hybrid promiscuous (Hypr) GGDEF enzyme, Bd0367, from the predatory bacterium *B. bacteriovorus* HD100. Hallberg *et al.* previously identified Bd0367 and showed *in vitro* that it functions as both a cGAMP synthase and a diguanylate cyclase (48). Additional studies with Bd0367 have discovered that this enzyme is essential for completing the host-dependent predatory life cycle and that each signaling molecule, c-di-GMP and cGAMP, regulates independent aspects of the predatory process (34). However, the molecular mechanism that causes Bd0367 to switch its production between c-di-GMP to cGAMP is still unknown.

Prior to constructing the dual metabolite sensor, the two RNA-based biosensors were analyzed individually for their binding affinity and selectivity under physiologically relevant conditions (3 mM Mg^2+^ and at 37 °C) within a tRNA scaffold to model their activity within the tRNA-DF30 scaffold (Figure 2A-B, Supplementary Figure S7). For c-di-GMP signaling detection, we used the Pl-B Pepper biosensor developed for this study, and for cGAMP sensing, a previous biosensor, Gm0970 Spinach (14), was modified with the Spinach2 aptamer to improve biosensor fluorescence activation in cells (49). The Pl-B Pepper c-di-GMP biosensor in tRNA scaffold exhibited nanomolar binding affinity (*K*_d_= 47 ± 8 nM) for c-di-GMP, while binding affinity to off-target ligand cGAMP was over 230-fold poorer (*K*_d_ = 11 ± 3 µM) (Figure 2A). The cGAMP biosensor, Gm0970 Spinach2, exhibited a binding affinity in the low micromolar range (*K*_d_= 1.3 ± 0.1 µM) for cGAMP, with binding affinity to off-target c-di-GMP outside the tested range of concentrations (*K*_d_ > 50 µM) (Figure 2B). These results confirm that both cGAMP and c-di-GMP biosensors selectively bind to their cognate ligands and therefore, the Pl-B and Gm0970 biosensors are compatible for dual metabolite sensor design, though the former is more sensitive than the latter.

**Figure 2.**
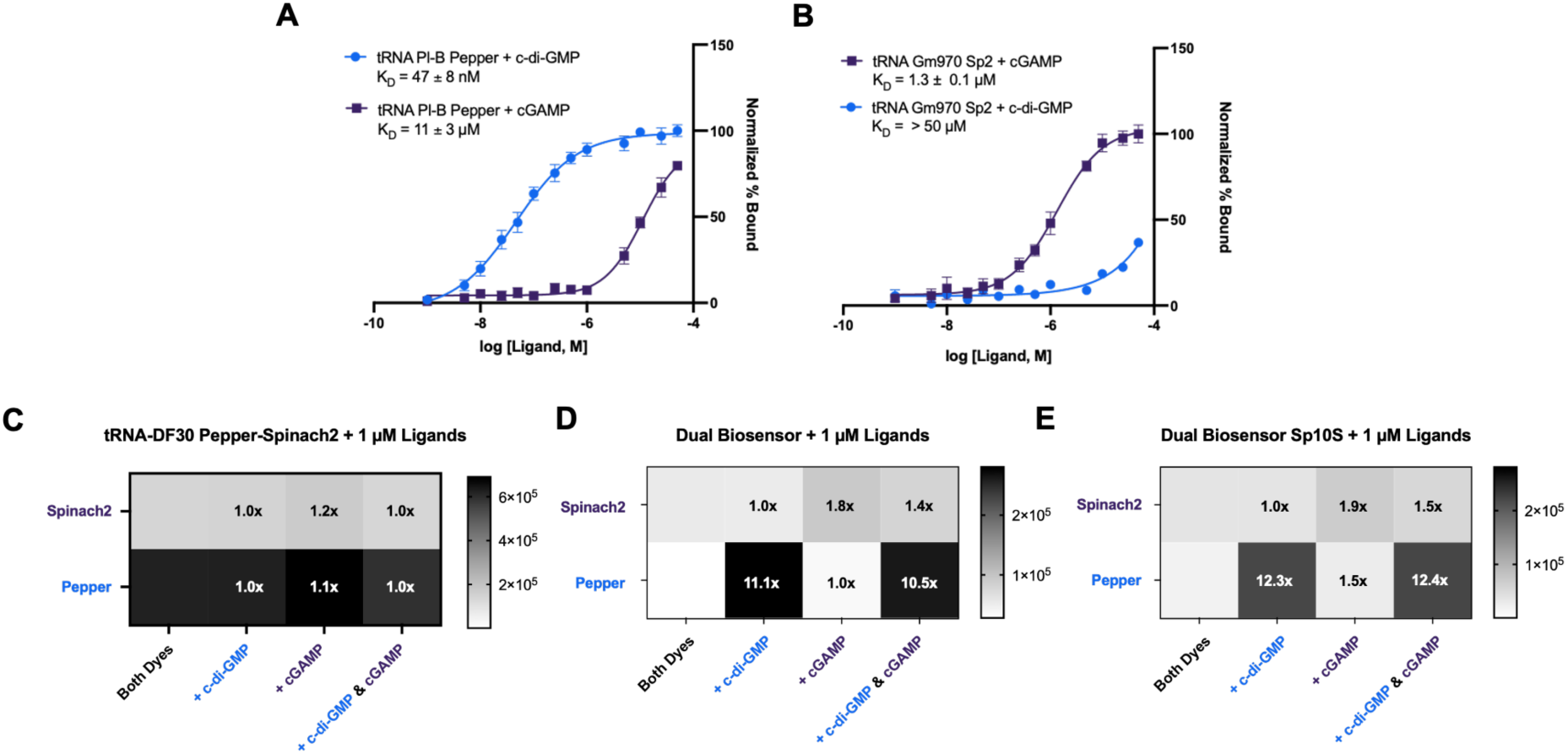
*In Vitro* Analysis of Individual RNA-based Biosensors and the Dual Biosensors. (A) Biosensor binding affinity measurements of tRNA scaffolded Pl-B Pepper are shown as normalized % bound (maximum fluorescence), normalized to Pl-B Pepper with its associated ligand target (c-di-GMP), with titration of c-di-GMP or 3’3’ cGAMP at 3 mM Mg2+, 37 °C. (B) Biosensor binding affinity of tRNA scaffolded Gm0970 Spinach2 is shown as normalized % bound (maximum fluorescence), normalized to Gm0970 Spinach2 with its associated ligand target (3’3’ cGAMP), with titration of c-di-GMP or 3’3’ cGAMP at 3 mM Mg2+, 37 °C. Points and error bars represent averages and standard deviations from three independent replicates for all binding affinity measurements. Best-fit curves are also shown, and the KDs of the biosensor affinity to ligands are listed in the legend. (C-E) Fluorescent activation of tRNA-DF30 Pepper-Spinach2 (control) and biosensor constructs with 1 µM of c-di-GMP and 3’3’ cGAMP, along with 10 µM DFHBI-1T and 200 nM HBC620. Fold activation is determined by looking at the fluorescent intensity of Spinach2 and Pepper aptamers with only DFBHI-1T and HBC620 present. The data shown are the average of three independent replicates.

Following this, the selectivity of the dual metabolite sensor was examined *in vitro* with two different dual metabolite sensor designs (Figure 2C-E, Figure 3A, Supplementary Figure S8, Supplementary Table S1). Both designs are based on the tRNA-DF30 Pepper-Spinach2 construct (47), which serves as a control and gives constitutive fluorescence in two colors. In the first construct, named Dual Biosensor, Pl-B Pepper is substituted in place of the Pepper aptamer, and Gm0970 Spinach2 (Sp2) in place of Spinach2. The second design, Dual Biosensor Sp10S, is identical except for the insertion of a 10 bp spacer between the F30 scaffold and the Gm0970 Sp2 arm. This spacer was included to minimize potential steric clashes or RNA misfolding by spatially separating the biosensor by almost one helical turn.

**Figure 3.**
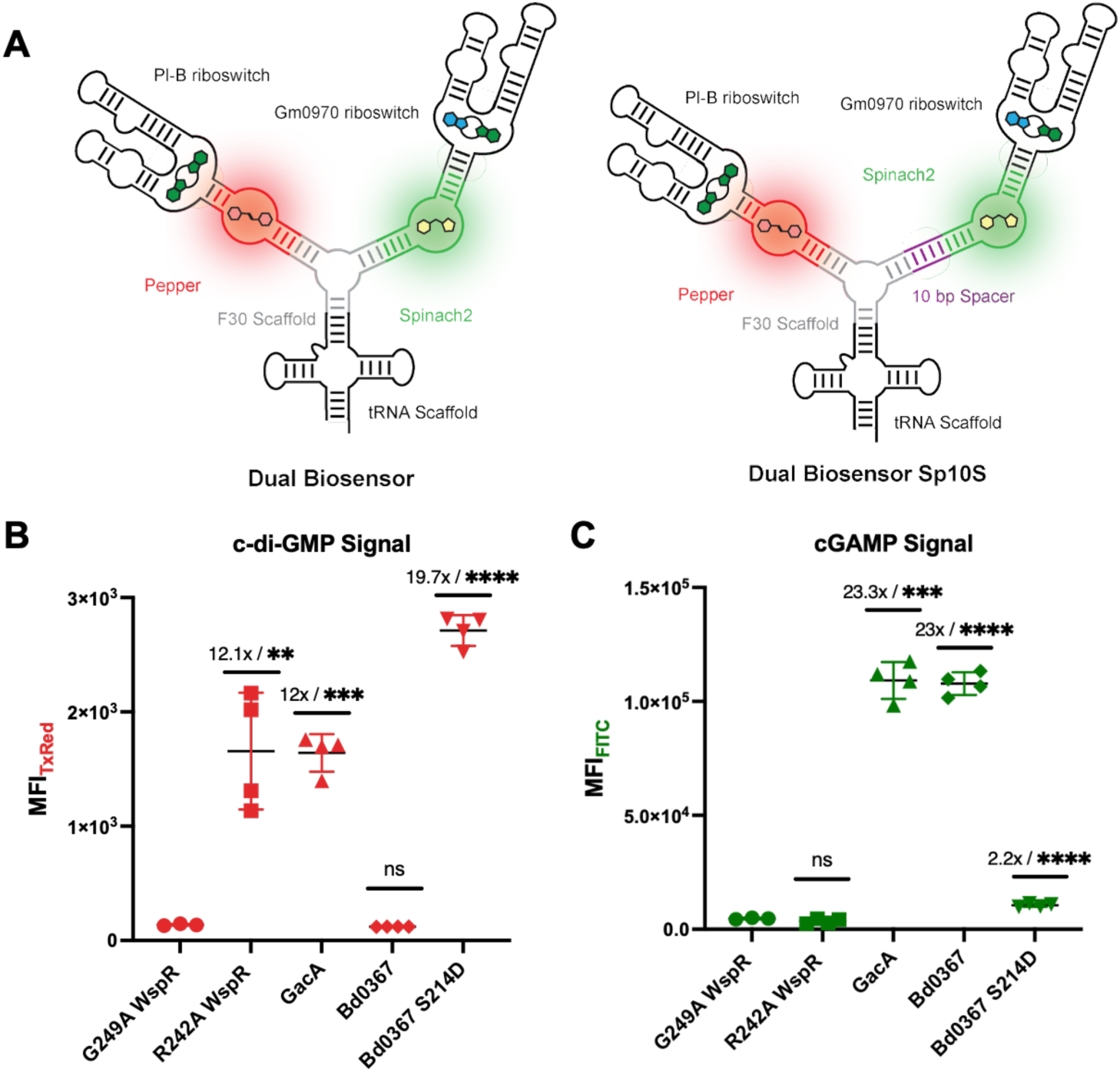
Characterization of Dual Biosensors Function and Sensitivity *In Vivo*. (A) Cartoon representation of Dual Biosensors. Both Dual Biosensor (left) and Dual Biosensor Sp10S (right) are within the tRNA-DF30 scaffold, where the tRNA scaffold (black) with fused to the three-way junction F30 scaffold (gray). Ligand binding of c-di-GMP Pl-B riboswitch enables the Pepper aptamer (red) to bind and activate the red HBC fluorogenic dye (HBC620). Ligand binding of cGAMP Gm0970 riboswitch enables the Spinach2 aptamer (green) to bind and activate the green fluorogenic DFHBI-1T dye. In purple is the 10 base-pair spacer that spatially moves the Gm0970 Spinach2 sensor on Dual Biosensor Sp10S by approximately one helical turn. (B-C) Raw MFI values in the TxRed Channel (B) and FITC Channel (C) of the Dual Biosensor Sp10S with various CDN synthase constructs. Data are from four biological replicates, by analyzing 30,000 cells per sample analyzed after 1 h incubation with DFHBI-1T (50 µM) and HBC620 (200 nM). Each point is the mean of the four biological replicates tested with each CDN synthase construct, and error bars are the standard deviation. Statistical comparison was performed by a two-tailed *t*-test. ns *P* > 0.05 / \*\**P* < 0.01/ *\*\*\*P* < 0.001/ \*\*\*\**P* < 0.0001.

Based on the binding affinity analyses in Figure 2A and B, 1 µM of each ligand was chosen for *in vitro* experiments, as each biosensor with 1 µM of its cognate ligand is approximately 50% or greater in its bound state (tRNA Pl-B Pepper + c-di-GMP: 89% bound; tRNA Gm0970 Sp2 + cGAMP: 48% bound). Furthermore, tRNA-DF30 Pepper-Spinach2 was tested to confirm that the cyclic dinucleotide ligands had negligible effects on the fluorogenic RNA aptamers themselves. The biosensor response for cyclic dinucleotides, c-di-GMP and cGAMP, can be determined at separate emission wavelengths due to the choice of fluorogenic dyes. For the c-di-GMP levels, Pepper-HBC620 signal is measured in the TxRed emission channel (Pepper-HBC620; ex/em 577/620), and for cGAMP levels, Spinach2-DFHBI-1T is measured in the FITC emission channel (Spinach2-DFHBI-1T; ex/em 482/505).

As expected, the tRNA-DF30 Pepper-Spinach2 control showed constitutive fluorescence and no modulation upon incubation with c-di-GMP or cGAMP (Figure 2C). For the Dual Biosensor, the addition of 1 µM cGAMP resulted in a 1.8-fold increase in fluorescence from the Gm0970-Spinach2 sensor, while the addition of 1 µM c-di-GMP resulted in an 11.1-fold increase with the Pl-B Pepper sensor (Figure 2D). Fold change values are reported relative to the background fluorescence observed with both dyes, DFHBI-1T and HBC620, incubated with the RNA but no ligands. Interestingly, moderate attenuation of the fluorescence signal was observed when both ligands are present simultaneously, likely due to subtle structural interference between sensors.

The Dual Biosensor Sp10S also showed selective fluorescence response from each sensor in response to its cognate ligand, with moderate signal attenuation when both ligands are present (Figure 2E). However, in this construct with the additional spacer, Pl-B Pepper appears to exhibit a low-level response (1.5-fold) to cGAMP, suggesting minor background crosstalk. Since the tRNA scaffold enhanced the *in vitro* binding affinity of the Pl-B Pepper biosensor, the tRNA-DF30+10S spacer may have enhanced the binding affinities even more than the other scaffolds, such that signal crosstalk is observed at 1 µM of cGAMP, which is not observed for the Dual Biosensor without a spacer.

Taken together, both Dual Biosensor constructs function as orthogonally fluorescent, ligand-selective sensors for the simultaneous detection of c-di-GMP and cGAMP. While the Pl-B Pepper-HBC620 arm consistently demonstrates more robust fluorescence activation, both individual biosensors appear to respond independently and specifically to their respective ligands. These results show the feasibility of a two-color dual metabolite RNA-based biosensor capable of resolving distinct cyclic dinucleotide signals with minimal crosstalk.

### Monitoring Cyclic Dinucleotide Signaling in Live Cells Using the Dual Metabolite Sensor

Next, the dual sensor construct was tested for its capability to detect c-di-GMP and cGAMP levels in live cells. Plasmids encoding Dual Biosensor Sp10S were co-expressed in *E. coli* with different enzymes known to alter intracellular levels of c-di-GMP or cGAMP, and cellular fluorescence was analyzed by flow cytometry (Figure 3). For this analysis, the enzymes used were the diguanylate cyclase WspR with an allosteric inhibitor site mutation (I-site, R242A) (50), which blocks feedback inhibition of c-di-GMP production, G249A WspR (51), a catalytically inactive WspR variant, and the Hypr GGDEF, GacA (48, 52), which predominantly makes cGAMP with c-di-GMP as a minor product. The raw median fluorescence (MFI) values from cells expressing active cyclic dinucleotide synthases were compared to the MFI values with the inactive G249A WspR mutant (Figure 3B-C). For c-di-GMP signal levels, cells expressing active WspR showed a 12.1-fold increase with Dual Biosensor Sp10S. For cGAMP levels, cells expressing wild-type GacA, exhibited a 12-fold activation with the Dual Biosensor Sp10S. These initial results demonstrate that this dual metabolite sensor design can distinguish increased production of c-di-GMP or cGAMP signals in cells by giving distinct fluorescence outputs.

Additionally, we tested the ability of the dual metabolite sensor to detect c-di-GMP and cGAMP levels when co-expressed with variants of Bd0367, our enzyme of interest. The Bd0367 enzyme primarily produces cGAMP, with c-di-GMP as a minor product (34). The I-site of Bd0367 was mutated (R260A) as well to prevent feedback inhibition that could decrease its activity; this I-site mutant was previously shown to still be capable of producing both c-di-GMP and cGAMP *in vitro* (34). The second Bd0367 variant analyzed is the S214D R260A mutant, as Ser214 is found in the active site of Hypr GGDEFs whereas Asp214 is found in canonical GGDEFs, so this mutation switches the enzyme to produce almost exclusively c-di-GMP (34). As expected, the I-site R260A Bd0367 mutant exhibited strong cGAMP signal fold activation (23-fold) compared to the inactive G249A WspR control, and appears to be comparable in activity to GacA. Conversely, cells expressing the S214D mutant showed a 19.7-fold increase in c-di-GMP signal and a 10-fold decrease in cGAMP signal, showing the complete switch in product selectivity

The raw MFI values of each synthase in the individual FITC and TxRed channels display a selective CDN signal output detection to their known function compared to the inactive G249A WspR mutant, except for GacA in the TxRed channel, as c-di-GMP is a minor synthase product (Figure 3B-C). Furthermore, as expected, the I-site R260A Bd0367 mutant exhibited strong cGAMP signal fold activation compared to the inactive G249A WspR control, and appears to be much more active than GacA. Conversely, cells expressing the S214D mutant showed a significant increase in the c-di-GMP signal, indicating a complete switch in product selectivity. These data show that the dual metabolite sensor can detect production levels of c-di-GMP and cGAMP in the presence of known cyclic dinucleotide synthase activities in live cells. Thus, the Dual Biosensor Sp10S design is a suitable construct for observing changes in CDN production of signaling molecules using the signal-switching Hypr GGDEF enzyme, Bd0367, from *B*. *bacteriovorus* in subsequent cell-based studies.

### Investigation of Histidine Kinase Candidates and Their Cognate Activity with Bd0367

Since the discovery that Bd0367 synthesizes both c-di-GMP and cGAMP to promote different stages of the predatory life cycle in *B. bacteriovorus*, the mechanism that regulates its switch between these two signals and the conditions under which this occurs remain intriguing. One possible molecular mechanism, since Bd0367 contains an N-terminal receiver (REC) domain with a canonical phosphorylation site (D74), is that it may be part of a two-component system. Bacterial two-component systems consist of a histidine kinase that senses external or internal stimuli and transfers a phosphate to the REC domain of a response regulator protein, which then triggers changes in cellular behavior. Furthermore, it has been observed *in vitro* that a D74E phosphomimic mutant has a slight product distribution shift in favor of c-di-GMP over cGAMP, although this mutant showed overall decreased activity (34). No histidine kinase has been identified yet to be associated with Bd0367; thus, the dual sensor was applied to examine the effect of candidate histidine kinases from *B*. *bacteriovorus* on Bd0367 activity and product switching.

We hypothesized that co-expression of the cognate histidine kinase might induce a shift from cGAMP to c-di-GMP production, or vice versa, potentially through phosphorylation of the REC domain or alternative activation mechanisms. To identify candidate histidine kinases (HKs) potentially involved with Bd0367, the *B*. *bacteriovorus* transposon sequencing (Tn-Seq) dataset from Duncan *et al* was analyzed (44). In this dataset, the authors report gene fitness scores (*W*) that reflect the survivability and fitness of *B. bacteriovorus* under various predatory conditions, including different prey species and growth states. Fitness scores of all annotated HKs were compared to the fitness scores from the Bd0367 knockout strain. Histidine kinases that exhibited fitness scores phenocopying or mirroring the significant differences observed in the Bd0367 knockout mutant were selected for further investigation, as similar fitness profiles under the tested conditions suggest a potential role in the same regulatory pathway (44). Using this approach, four uncharacterized histidine kinases, Bd3648, Bd0596, Bd1382, and Bd1828, were initially identified as suitable candidates. However, the initial analysis misaligned the *B. bacteriovorus* 109J locus tags in the Tn-Seq dataset (44) to KEGG database annotations (53). After correcting this issue, Bd0596 remained the top candidate, whereas Bd3648, Bd1382, and Bd1828 are less likely to be relevant HK candidates (Supplementary Table S2).

The genes encoding HK candidates Bd0596 and Bd3648 were codon optimized for *E. coli* and successfully cloned into dual expression plasmids also containing Bd0367 R260A. Bd0367 activity was then assessed by measuring c-di-GMP and cGAMP levels through flow cytometry with the dual metabolite sensor within a second plasmid (Figure 4). The fluorescence response for each biosensor is observed as the ratio of median fluorescence intensities (MFIs) in the FITC (cGAMP sensor) and TxRed (c-di-GMP sensor) emission channels. The proportional relationship between the FITC and TxRed channels serves as a ratiometric readout of cellular levels of each cyclic dinucleotide that internally corrects for cell-to-cell variability in expression levels and can be normalized to known enzyme controls. Thus, the TxRed / FITC ratio shows increases in c-di-GMP relative to cGAMP, whereas the inverse, FITC / TxRed ratio, shows increases in cGAMP relative to c-di-GMP. For clarity, both ratios are shown. Cells co-expressing HK Bd3648 and Bd0367 R260A did not significantly alter the cyclic dinucleotide enzymatic production ratio compared to Bd0367 R260A alone (Figure 4B-C). In contrast, co-expression with HK Bd0596 resulted in a marked shift in product distribution, favoring c-di-GMP over cGAMP to a similar extent to the Bd0367 S214D mutant. These results demonstrate that the dual metabolite sensor is a sensitive tool capable of detecting changes in c-di-GMP and cGAMP product ratios in response to HK co-expression.

**Figure 4.**
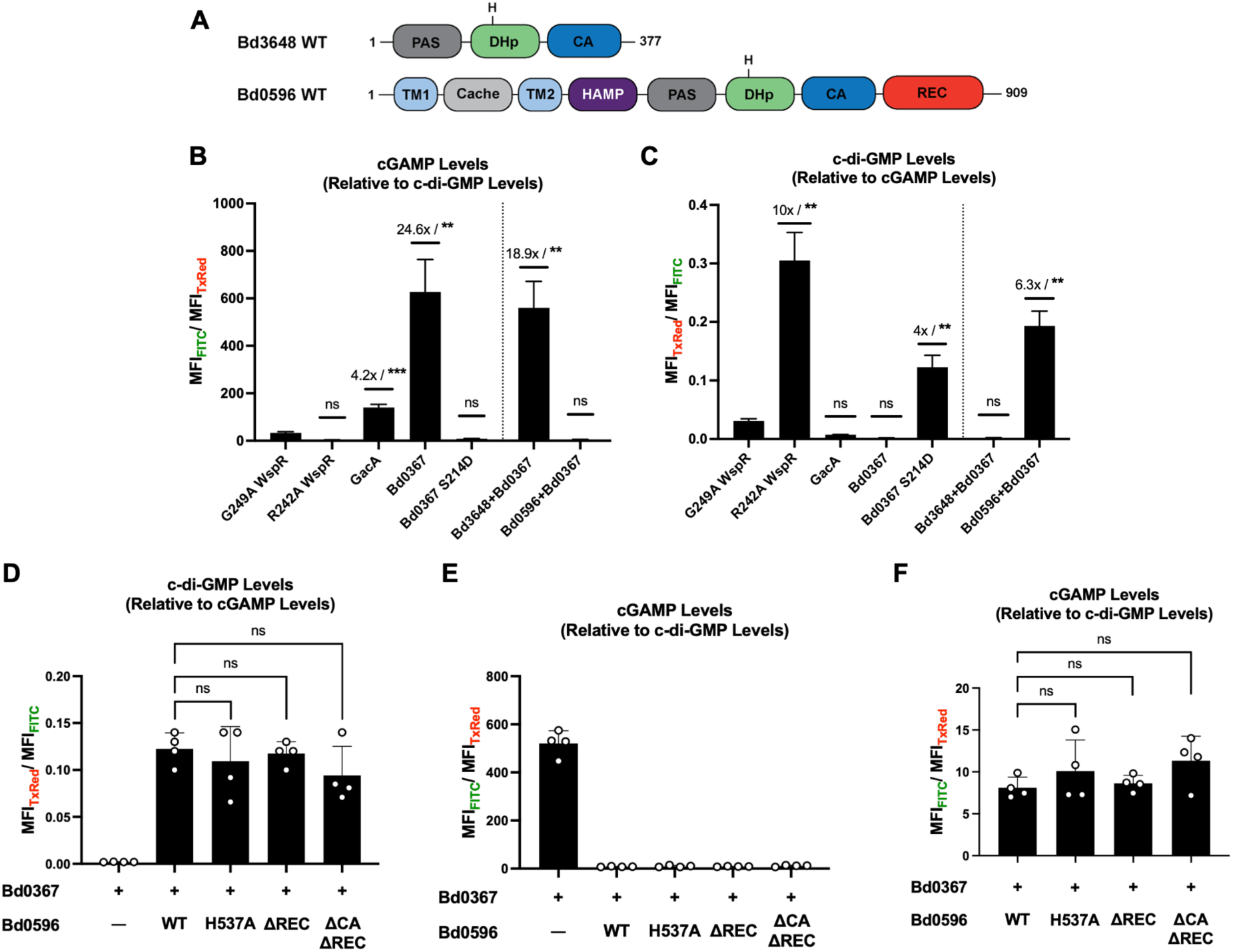
Investigating Histidine Kinase Candidates with the Dual Biosensor. (A) Domain organization of the tested histidine kinase candidates, Bd3648 and Bd0596. TM1 and TM2 stand for transmembrane domains, DHp for the dimerization and histidine phosphotransfer domain, CA for the ATP binding domain, and REC for the receiver domain. (B-C) Median fluorescence intensities (MFI) are measured via flow cytometry with E. coli BL21 (DE3) Star cells co-expressing Dual Biosensor Sp10S with various cyclic dinucleotide synthases. MFI values of FITC and TxRed channels are normalized to each other to observe the relative c-di-GMP and cGAMP levels within cells. For c-di-GMP Levels, this is represented as MFI-TxRed/ MFI-FITC, and for cGAMP, this is MFI-FITC/ MFI-TxRed. Fold activation is determined by the mean production ratio value of inactive G249A WspR compared to active CDN synthases. Data are from three or four biological replicates by analyzing 30,000 cells per sample after 1 h incubation with DFHBI-1T (50 µM) and HBC620 dye (200 nM). Data represented the mean with a standard deviation of three or four production ratios. (D-F) Median fluorescence intensities (MFI) are measured via flow cytometry with E. coli BL21 (DE3) Star cells co-expressing Dual Biosensor Sp10S with various Bd0596 histidine kinase mutants. MFI values of FITC and TxRed channels is normalized to each other to observe the relative c-di-GMP and cGAMP levels within cells. For c-di-GMP Levels, this is represented as MFI-TxRed/ MFI-FITC, and for cGAMP this is MFI-FITC/ MFI-TxRed. (D) c-di-GMP levels relative to cGAMP levels and (E-F) cGAMP levels relative to c-di-GMP levels, where (E) is the full dataset and (F) is a close-up on Bd0596 mutants. Data are from four biological replicates by analyzing 50,000 cells per sample after 1 h incubation with DFHBI-1T (50 µM) and HBC620 dye (200 nM). Data represents the mean with the standard deviation of four production ratios. Statistical comparison was performed by a two-tailed *t*-test. ns *P* > 0.05 / \*\**P* < 0.01/ *\*\*\*P* < 0.001/ \*\*\*\**P* < 0.0001.

### Probing the Relationship Between Histidine Kinase Bd0596 and Rec-GGDEF Enzyme Bd0367

To evaluate HK Bd0596 activity and its effect on Bd0367 product ratios, a series of Bd0596 mutants were constructed and tested in cells based on their predicted domains (Supplementary Figure S9). The mutants contain an alanine point mutation at the histidine phosphorylation site (H537), which is critical for HK catalytic activity, or deletions of the C-terminal REC and ATP-binding kinase (CA) domains (54–56). Preliminary results with four biological replicates for each Bd0596 construct unexpectedly revealed that the loss of HK catalytic activity or REC/CA deletions did not significantly change its effect on Bd0367 activity (Figure 4D-F).

The above results suggest that the observed large shift toward c-di-GMP production by Hypr GGDEF Bd0367 is not due to the HK activity of Bd0596. An explanation we considered is that overexpression of the membrane-bound HK is masking a more specific interaction by non-specifically disrupting bacterial membrane integrity. This disruption can lead to depletion of cellular ATP, which is a substrate for cGAMP but not c-di-GMP synthesis (57–60). To test this, the ATPase inhibitor carbonyl cyanide m-chlorophenyl hydrazone (CCCP) (42) was used to assess whether Bd0367 activity is substrate-limited under ATP-depleting conditions (Supplementary Figure S10). Flow cytometry analysis showed a significant increase in intracellular c-di-GMP levels, as evidenced by relative ratios and raw MFI values. This result indicates that the full-length, membrane-bound HK Bd0596 may have skewed the cellular and enzymatic activity profile when co-expressed with Bd0367.

To eliminate the ATP-depletion artifact caused by overexpression of a membrane-bound HK, another set of experiments was performed with cytosolic versions of Bd0596 (ΔTM), which lack the predicted transmembrane domains. These studies were conducted with a sample size of n=20 to increase the statistical power of the dataset, as the cytosolic constructs are not expected to be as active. WT was tested alongside phosphorelay mutants with H537A to knock out the canonical histidine and/or deletion of the C-terminal REC domain. In the initial experiment, no significant change in product ratios was observed comparing WT to the H537A mutant, although a slight trend was observed (Supplementary Figure S11). In a more extensive analysis of cytosolic Bd0596 variants, a significant change in product ratios was observed between the active Bd0596ΔTM and the phosphorelay mutants (Figure 5A-B). Specifically, the active Bd0596ΔTM construct appears to modestly reduce c-di-GMP and increase cGAMP production by Bd0367 compared to the three inactive phosphorelay mutants.

**Figure 5.**
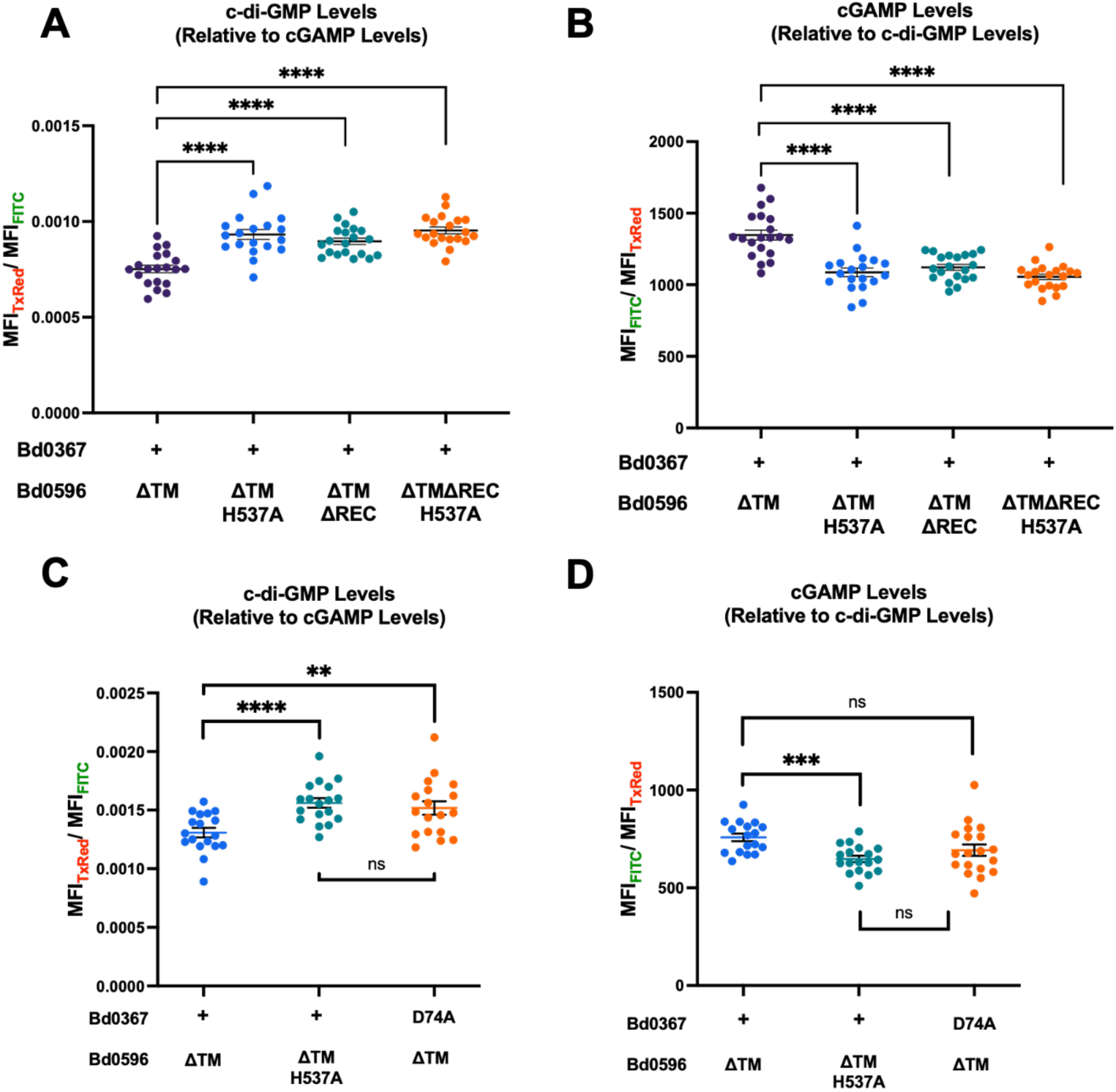
Cytosolic Bd0596 Mutation and Phosphotransferase Mechanism Analysis on CDN Levels in Cells. Median fluorescence intensities (MFI) are measured via flow cytometry with E. coli BL21 (DE3) Star cells co-expressing Dual Biosensor Sp10S with cytosolic Bd0596 histidine kinase mutants or with D74A Bd0367. MFI values of FITC and TxRed channels are normalized to each other to observe the relative (A) Analysis of cytosolic Bd0596 mutants. In the top left panel are the relative c-di-GMP levels, and in the top right panel are the relative cGAMP levels. (B) Analysis of the phosphotransferase mechanism between Bd0596 and Bd0367. In the bottom left panel are the relative c-di-GMP levels, and in the bottom right panel are the relative cGAMP levels. Data are from 20 biological replicates by analyzing 50,000 cells per sample after 1 h incubation with DFHBI-1T (50 µM) and HBC620 dye (200 nM). Outliers were identified and excluded from the data using the ROUT method; Q=5 %. Statistical comparison was performed by a two-tailed *t*-test where n=17, 18, 19, or 20. ns *P* > 0.05/ \**P* ≤ 0.05/\*\**P* < 0.01/ *\*\*\*P* < 0.001/ \*\*\*\**P* < 0.0001. Data represent the mean ± SEM.

We next examined the role of the conserved aspartate residue (D74) in the N-terminal REC domain in Bd0367, which is the canonical phosphorylation site in two-component systems. Prior work showed that the phosphomimetic D74E mutation modestly shifts Bd0367 product distribution toward c-di-GMP *in vitro* (34). However, the functional impact of a non-phosphorylatable D74A variant has not yet been assessed. Encouragingly, the effect on product ratios of knocking out the canonical REC phosphorylation site with Bd0367 D74A is the same as knocking out the canonical HK histidine with Bd0596 H537A (Figure 5C-D). The lack of an additive effect between the two critical phosphotransfer residues supports the model in which Bd0596 and Bd0367 participate in the same signaling pathways, potentially through a phosphotransfer from the HK to the D74 residue in the REC domain of Bd0367.

Taken together, these results provide some evidence that Bd0596 is the cognate HK that regulates the activity of Hypr GGDEF Bd0367 through phosphorylation. *In vitro* data on the D74E phosphomimic mutant of Bd0367 initially led us to hypothesize that phosphorylation would induce a shift to favor c-di-GMP production. Overexpression of full-length, membrane-bound Bd0596 supported this expectation, but more careful experiments incorporating mutant controls revealed that the product ratio shifts observed for full-length Bd0596 were independent of its HK activity, and likely were pleiotropic effects. Instead, with cytosolic Bd0596ΔTM constructs, it was found that the active HK consistently favors Bd0367 production of cGAMP, and this effect requires the conserved active site histidine H537 and C-terminal REC domain for the hybrid histidine kinase, as well as the conserved D74 phosphorylation site of the REC domain. Potential reasons why the observed modulation of cyclic dinucleotide product ratios is modest include (a) the cytosolic version of HK Bd0596 is weakly active due to being mislocalized and missing the extracellular sensor and transmembrane domains that regulate dimerization; (b) under cellular overexpression conditions, Hypr GGDEF Bd0367 already favors cGAMP production.

## DISCUSSION

This study demonstrates the feasibility of converting a previously published second-generation c-di-GMP Pl-B Spinach2 biosensor for c-di-GMP detection to incorporate the Pepper aptamer. Both Pl-B Pepper and Spinach2 sensors can effectively detect intracellular c-di-GMP concentrations in *E. coli*. Notably, Pl-B Pepper (**TS1.1**) exhibited a robust 37.5-fold fluorescence increase with wild-type WspR, whereas Pl-B Spinach2 only showed a 4-fold activation with overactive WspR D70E. Collectively, these results validate Pl-B Pepper as a sensitive and reliable alternative to Pl-B Spinach2 for intracellular c-di-GMP detection and highlight its suitability for use in a dual-metabolite RNA-based biosensor.

Replacing Spinach2 with the Pepper aptamer altered the sensor’s properties, including binding affinity, ligand specificity, and cellular brightness. Under similar conditions (3 mM Mg^2+^ and at 37 °C), Pl-B Pepper exhibited a dissociation constant (*K*_d_) for c-di-GMP of ∼215 nM, compared to Pl-B Spinach2 (*K*_d_ of ∼12 nM). However, this reduced affinity was accompanied by improved selectivity, with approximately 15% lower off-target activation to the off-target linear cleavage product of c-di-GMP, pGpG. While this reduced affinity may limit its sensitivity for detecting low or endogenous c-di-GMP levels when CDN synthases are not overexpressed, this can be mitigated with an alternative sensor design. Pl-B Pepper sensor variants (**TS2** and **TS2.1**) provide a promising alternative with a tighter binding affinity to c-di-GMP (20nM), depending on experimental needs. Additionally, the implementation of RNA scaffolds can increase sensor sensitivity, as observed with tRNA-scaffolded Pl-B Pepper, which exhibits a dissociation constant (K*_d_*) of ∼47 nM, reflecting a 4.5-fold improvement in c-di-GMP binding affinity compared to the non-scaffolded sensor.

A practical consideration for effective use of the Pepper-based sensor is that it requires pre-incubation with the HBC dye for approximately 1 h before fluorescent measurements, which may constrain experimental studies probing rapid dynamics. Despite this potential constraint, replacing the Spinach2 fluorogenic aptamer with Pepper provides spectral separation from existing green-fluorescent sensors, enabling dual-color detection systems (22, 25, 42, 50). Moreover, the chameleon-like ability of Pepper to fluoresce across a spectrum from cyan to red, depending on the HBC dye analog, allows Pepper-based sensors and tags to be readily used in combination with other fluorescent RNA-based sensors or protein tools in cells. Taken together, this combination of improved intracellular brightness and specificity makes Pl-B Pepper a sensitive and versatile component for the development of a dual biosensor.

Integration of two orthogonal biosensors, Pl-B Pepper for c-di-GMP and Gm0970 Spinach2 for cGAMP, into a single RNA transcript allowed for the development of a dual metabolite sensor with distinct color outputs for live-cell quantification of the two bacterial signaling molecules. This dual biosensor is distinct from other RNA-based ratiometric sensors, which usually detect a single metabolite in one channel and employ the other channel to normalize expression (9, 61–64). Although recent RNA-based biosensors have demonstrated the ability to sense multiple analytes *in vitro* and in cells, these systems have primarily focused on detecting biologically unrelated metabolites or transcriptional circuits composed of separate sensing components (61, 65, 66). In contrast, our dual biosensor detects two structurally similar and biologically relevant metabolites involved in bacterial signaling. Therefore, the development of this dual metabolite RNA-based sensor required stringent selectivity between the two biosensor components. Despite a single nucleotide difference between the cyclic dinucleotide signals cGAMP and c-di-GMP, these molecules regulate distinct signaling pathways. For instance, the Hypr GGDEF enzyme Bd0367 from *B. bacteriovorus* appears to switch from making c-di-GMP to cGAMP molecules at different stages of its predatory lifecycle (34, 67). Studies of Bd0367 have demonstrated that its production of c-di-GMP facilitates the swimming/attack phase, whereas after *B. bacteriovorus* consumes the intracellular nutrients of its prey, production of cGAMP by Bd0367 is required for productive prey escape (68).

Aside from the unique dual synthase activity of Bd0367, c-di-GMP and cGAMP play critical roles in other bacterial signaling pathways and in regulating multiple bacterial phenotypes. Cyclic di-GMP produced by diguanylate cyclase enzymes (GGDEF proteins) has been shown to regulate key cellular processes, including biofilm formation, host colonization, and bacterial virulence (69, 70). For cGAMP, it was first identified in *Vibrio cholerae*, where it contributes to virulence and intestinal colonization in mice, and is produced by the non-Hypr GGDEF cGAMP synthase DncV (71). In addition, cGAMP has also been implicated in an innate immune-like response in non-pathogenic bacteria, such as through the bacterial CBASS system, where detection of bacteriophage activates cGAMP synthesis; the resulting cGAMP then triggers *E. coli* cell death to prevent phage replication (72–74). Thus, the ability to monitor these bacterial signaling molecules in the same cellular context offers a powerful approach to dissect complex regulatory networks, microbial defense systems, and host-pathogen interactions.

To understand the mechanism underlying the production shift between c-di-GMP and cGAMP in Bd0367, we used the Dual Biosensor Sp10 sensor to dissect its enzymatic mechanism. Given that histidine kinases canonically activate response regulators (RRs) through transphosphorylation from a conserved histidine, including hybrid HKs, which are multidomain proteins containing a histidine kinase and a receiver (REC) domain within a single polypeptide. This domain architecture enables multi-step phosphorelays and introduces additional regulatory checkpoints and complexity into bacterial signaling pathways (60). However, recent studies have revealed an alternative activation pathway mechanism. For example, some HKs have been shown to allosterically activate their cognate RRs even in the absence of a phosphorylated histidine, thereby regulating RR activity through protein-protein interactions (75). Other activation mechanisms include cis-autophosphorylation, genetic bypass of essential HK-RR signaling, and modulation through HK sequestration with non-cognate RRs partners (56, 76–78).

Co-expression studies with histidine kinases (Bd0596 and Bd3648) in *E. coli* indicate that HK Bd0596 modulates the cyclic dinucleotide product ratio of Bd0367. Mutational studies support the regulatory partnership between Bd0596 and Bd0367 by testing the enzymatic phosphotransfer relay and the protein-protein interactions that drive signal transduction. Although co-expression of full-length Bd0596 mutants with Bd0367 produced large but likely pleiotropic changes in CDN levels, potentially due to the transmembrane domains, analysis of the cytosolic Bd0596 variants provided a cleaner mechanistic dissection of Bd0367 regulation. In particular, significant signal changes were observed between Bd0596 ΔTM and mutants that disrupted the phosphotransfer mechanism by mutating either of the two critical phosphotransfer residues, H537A in Bd0596 and D74A in Bd0367, suggesting that there is a functional link between Bd0596 and Bd0367.

While our data strongly indicate that Bd0596 is a regulatory partner of Bd0367, further experiments are needed to confirm the interaction and elucidate the precise mechanism of regulation. The absence of a perfect correlation indicates additional factors, such as multiple phosphorylation states, a direct protein-protein interaction between Bd0596 and Bd0367, or broader signaling networks, may modulate Bd0367 activity. Direct biochemical assays, such as *in vitro* phosphotransfer experiments, protein-protein interaction studies, and using phosphomimetic Bd0367 mutants, will clarify whether Bd0596 phosphorylates Bd0367. Additionally, applying the dual metabolite sensor in *B. bacteriovorus* rather than a heterologous *E. coli* system can offer deeper insights into the physiological relevance of the regulatory switch and native Bd0367 activity. After correctly linking the *B. bacteriovorus* 109J locus tag annotations from the Tn-Seq dataset to KEGG database annotations, other HK candidates that correlate well with the Bd0367 knockout strain are Bd0499, Bd3750, and Bd0584. Taken together, these future experiments will be essential to confirm the proposed Bd0596-Bd0367 regulatory pathway.

In summary, we developed an RNA-based dual metabolite sensor that utilized two orthogonal fluorogenic RNA-based biosensors, Pl-B Pepper and Gm0970 Spinach2, to dissect intracellular signaling pathways within bacteria. These RNA-based biosensors were chosen as c-di-GMP and cGAMP levels can be observed by a red (Pepper-HBC620) and green (Spinach2-DFHBI-1T) fluorescence output, respectively. Its ratiometric and orthogonal design enables multiplex monitoring of bacterial signaling within a single construct, offering a significant advantage over protein-based biosensors, which require co-expression of each sensor separately. The application of this dual metabolite sensor to analyze the activities of Hypr GGDEF Bd0367 and histidine kinase Bd0596 in live cells highlights the potential of RNA-based biosensors to uncover new signaling and regulatory pathway mechanisms.

## AUTHOR CONTRIBUTIONS

Madeline M. Mumbleau: Conceptualization, Formal analysis, Methodology, Validation, Writing—original draft. Ming C. Hammond: Conceptualization, Visualization, Writing—review & editing.

## ACKNOWLEDGEMENTS

The authors would like to thank Z. Hallberg for advice on data analysis of the *B. bacteriovoru*s Tn-Seq dataset and helpful discussions.

## FUNDING

This work was supported by grants from the National Science Foundation (1815508 to M.C.H.) and the National Institutes of Health (GM124589 to M.C.H.). The flow cytometry core facility is supported by the National Institutes of Health Award S10OD026959.

## CONFLICT OF INTEREST

The authors declare no competing financial interest.

## Notes

### Competing Interest Statement

The authors have declared no competing interest.

