## Supplementary Data for "Development of a Fluorescent RNA Biosensor for Dual Detection of cGAMP and c-di-GMP Signals in Live Bacteria"

### for

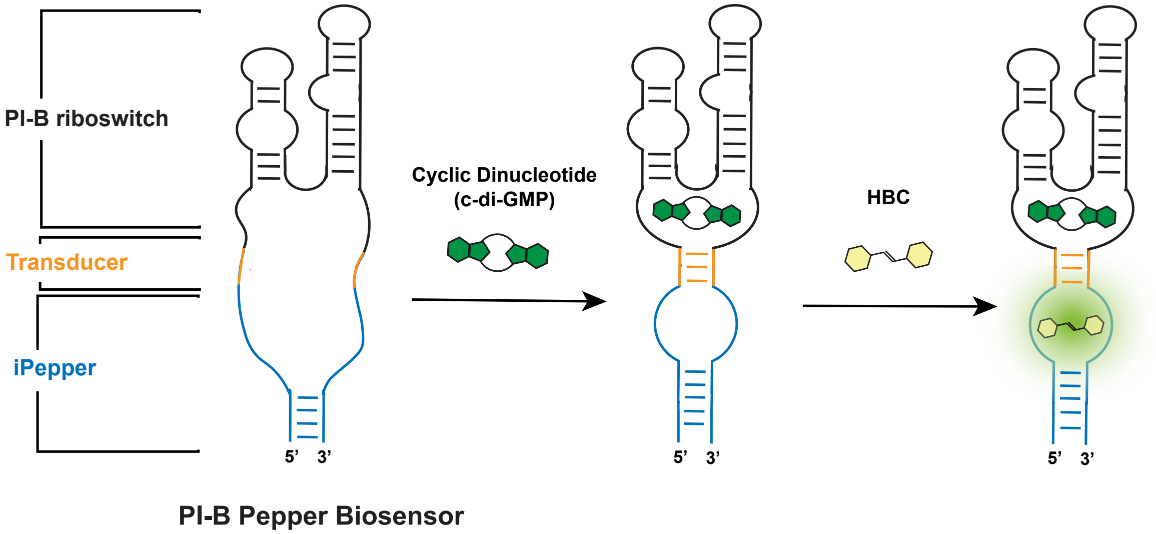


**Supplementary Figure 1. RNA-based Fluorescent Biosensors Detect Cyclic Dinucleotide Secondary Messengers.** Design scheme for a fluorescent biosensor that detects cyclic dinucleotide, c-di-GMP. Cyclic di-GMP binding to the PI-B riboswitch (black) enables the Pepper aptamer (green) to bind and activate a fluorogenic HBC dye molecule upon stabilization of the transducer domain (orange).

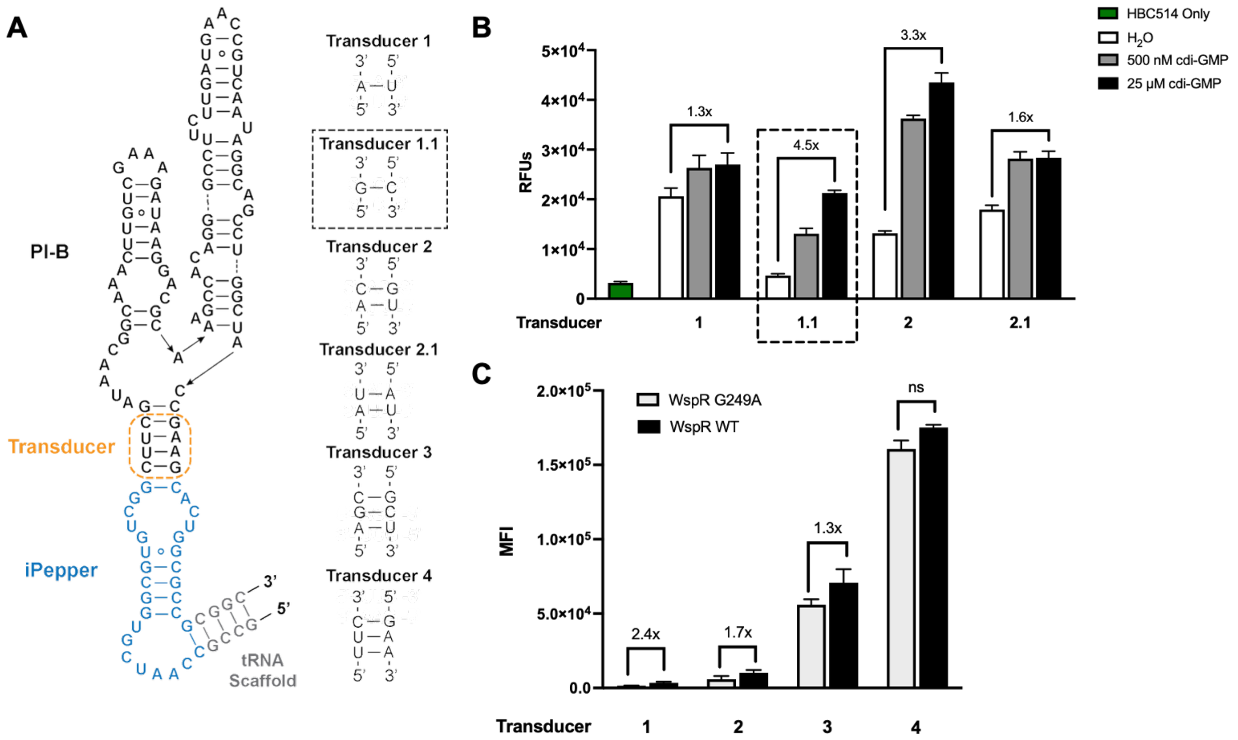


**Supplementary Figure 2. Initial PI-B Pepper Sensor Optimization and Screening *In Vivo*.** (A) Sequence and secondary structure model of PI-B Pepper construct, which comprises the PI-B riboswitch (black), a transducer domain region (boxed in orange), and the iPepper aptamer (blue). For *in vivo* studies, PI-B Pepper sensors are within a tRNA scaffold (gray). A series of transducer domains was tested to optimize fluorescence response by decreasing the fluorescence background signal. (B) Fluorescence activation of PI-B Pepper sensors and transducer domains in vitro with 0, 500 nM, and 25 μM c-di-GMP and measured as relative fluorescence units (RFUs). HBC514 dye background fluorescence is represented as a green bar. Fluorescence activation levels are based on the fold difference of fluorescence levels with 0 μM c-di-GMP versus 25 μM c-di-GMP. The optimal sensor is indicated with a black dotted box. The data shown are the average with a standard deviation of three independent replicates. (C) Average median fluorescence (MFI) measured by flow cytometry of *E. coli* BL21 (DE3) Star cells co-expressing biosensor constructs with different transducer domains as indicated above, with various diguanylate cyclase enzymes. Black represents an inactive mutant of diguanylate cyclase WspR (G249A); white represents wild-type WspR. Fluorescence activation is determined with the mean fluorescence of each PI-B Pepper construct under different enzyme conditions. The optimal sensor is indicated by a black dotted box. Data are from three biological replicates by analyzing 30,000 cells per sample after 1 h incubation with HBC514 dye (200 nM). Data are the average with a standard deviation of three biological replicates.

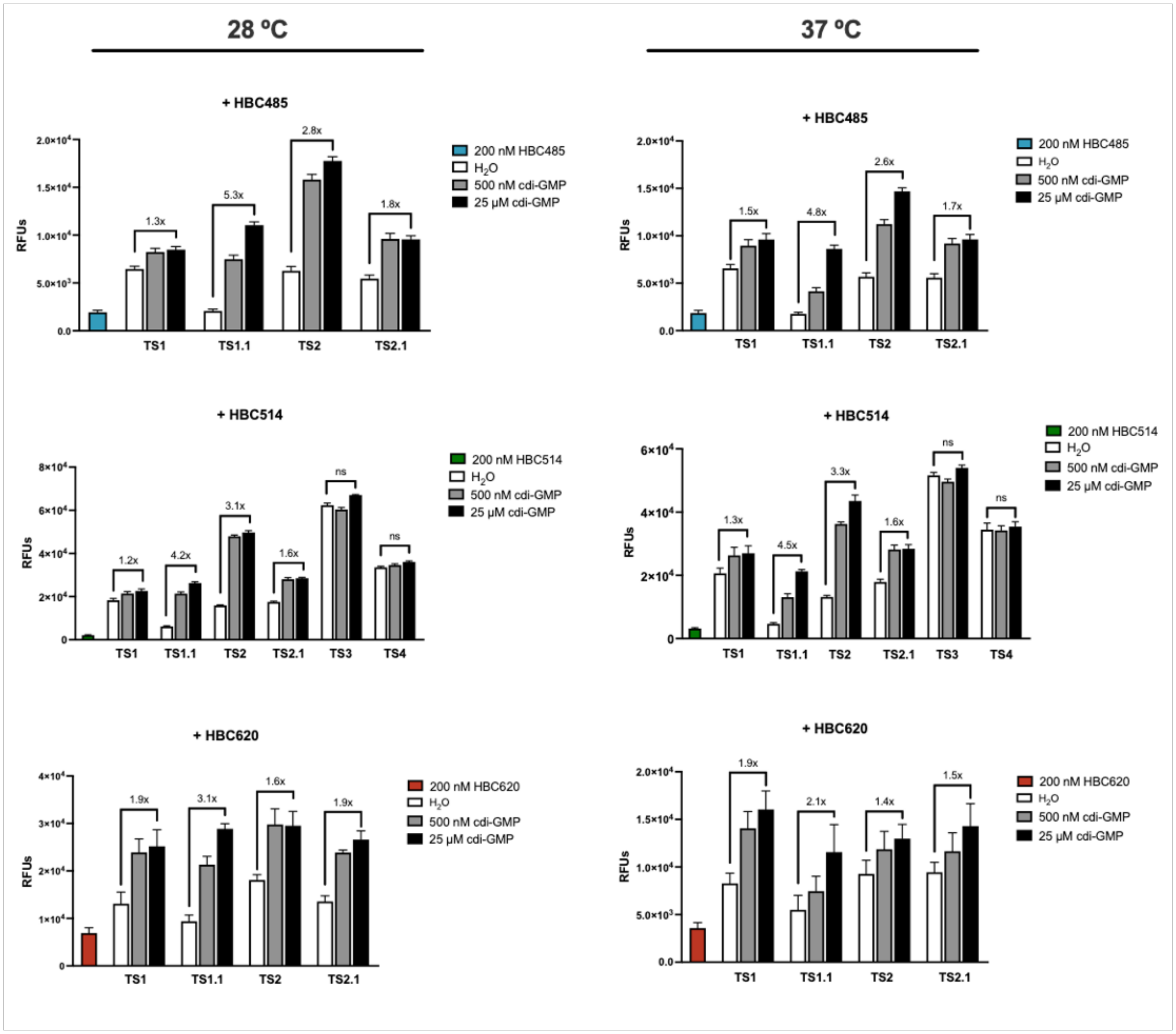


**Supplementary Figure 3. Fluorescent activation of PI-B Pepper in vitro with 200 nM HBC dyes, HBC485, HBC514, and HBC620.** PI-B Pepper with various transducer domains shown above is tested with different concentrations of c-di-GMP measured in RFUs with 3mM Mg2+ and 200nM HBC dyes (left panel) at 28 ºC and (right panel) at 37 ºC. HBC dye fluorescence backgrounds are represented as a blue (HBC485), green (HBC514), or red (HBC620) bar. Fluorescence activation levels are based on the fold difference of fluorescence levels with 0 μM c-di-GMP versus 25 μM c-di-GMP. The data shown are the averages with a standard deviation of three independent replicates.

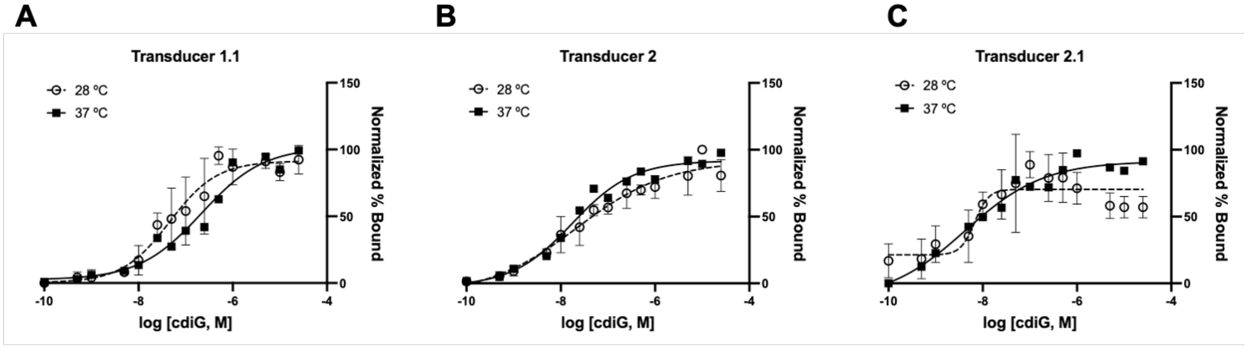


**D**

| Temp | Transducer 1.1 | Transducer 2 | Transducer 2.1 |
|---|---|---|---|
| 28 ºC | 46 ± 4 | ≤ 20 | ≤ 20 |
| 37 ºC | 215 ± 78 | ≤ 20 | ≤ 20 |

**Supplementary Figure 4. Biosensor binding affinities with a titration of c-di-GMP.** Titration of c-di-GMP with PI-B Pepper biosensors and HBC514 at varying temperatures (28 ºC –empty circles; 37 ºC – filled circles) with 3 mM Mg2+ at 37 ºC. Each panel displays a different PI-B Pepper biosensor (A. Transducer 1b, B. Transducer 2a, C. Transducer 2b). Points and error bars represent averages and standard deviation of three independent replicates, except for the 28 ºC data of Transducer 1b with two independent replicates. Best-fit curves are also shown (28 ºC – dotted lines; 37 ºC – solid lines). All binding affinity measurements included 100 nM HBC514 and 20 nM RNA. A table of KDs values in nM is displayed in D.

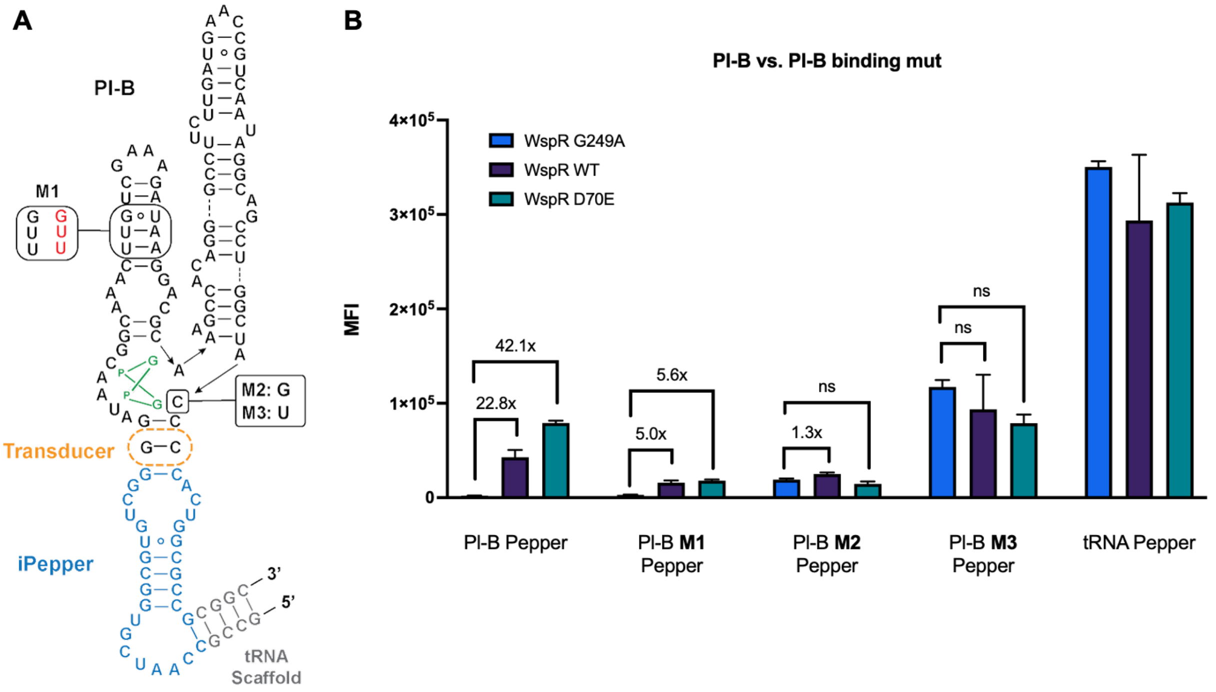


**Supplementary Figure 5. *In Vivo* Analysis of the Fluorescence Activation of PI-B Pepper Biosensor from a c-di-GMP Binding Event.** (A) Sequence and secondary structure model of PI-B Pepper sensor with optimal transducer domain 1b. Mutants analyzed in the study are boxed; c-di-GMP is shown in green. (B) Median fluorescence intensity (MFI) is measured by co-expressing PI-B Pepper or tRNA Pepper in *E. coli* BL21 (DE3) Star cells with various enzymes. Blue represents an inactive mutant of WspR (G249A); purple represents wild-type WspR; teal represents a constitutively active mutant of WspR (D70E). Fluorescence activation is indicated above by calculating the fluorescence fold difference of wild-type WspR or WspR D70E levels compared to WspR G249A. Data are from three biological replicates by analyzing 30,000 cells per sample after 1 h incubation with HBC514 dye (200 nM). Data are the average with a standard deviation of three biological replicates.

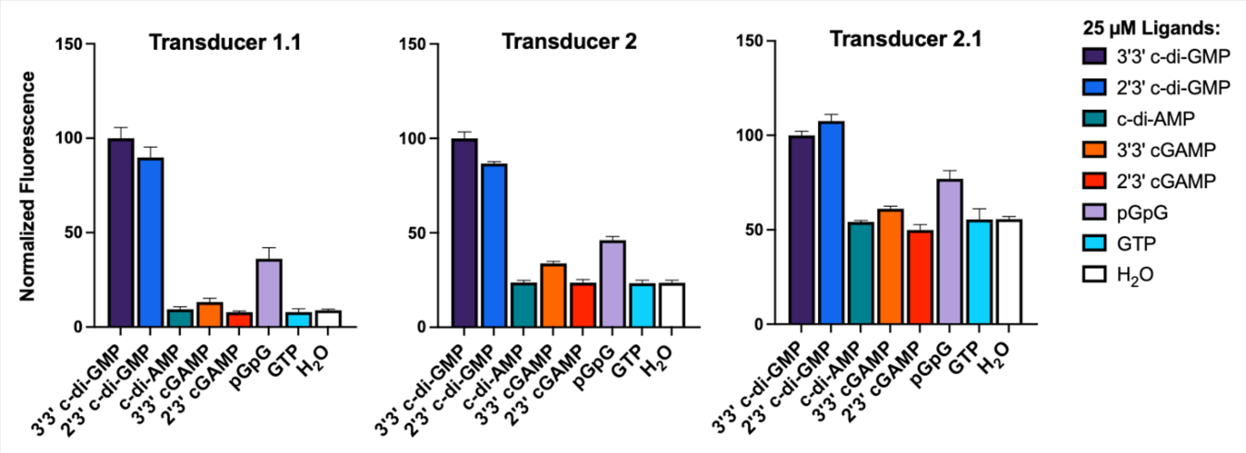


**Supplementary Figure 6. *In Vitro* Ligand Specificity of PI-B Pepper Sensors.** Biosensor ligand specificity of three PI-B Pepper sensors with ligands at 25 µM concentrations at 3 mM Mg2+, 37 ºC: 3’3’ c-di-GMP (purple), 2’3’ c-di-GMP (blue), c-di-AMP (green), 3’3’ cGAMP (orange), 2’3’ cGAMP (red), pGpG (light purple), GTP (light blue), and no ligand (white). For each biosensor, fluorescence levels were normalized to signal response with 3’3’ c-di-GMP.

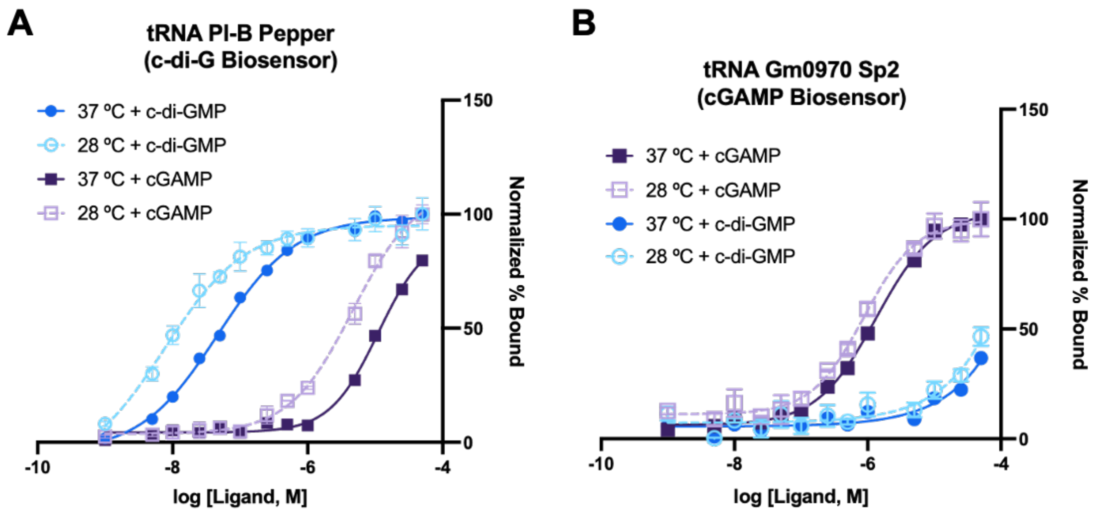


**C**

| Temp | tRNA PI-B Pepper | | tRNA Gm0970 Sp2 | |
|---|---|---|---|---|
| | **+ c-di-GMP** | + cGAMP | + c-di-GMP | **+ cGAMP** |
| 28 ºC | **≤ 5 nM** | 4.6 ±1 μM | > 50 μM | **870 ± 180 nM** |
| 37 ºC | **47 ± 8 nM** | 11 ± 3 μM | > 50 μM | **1.3 ± 0.1 μM** |

**Supplementary Figure 7. Full *In Vitro* Binding Curve Analysis of tRNA scaffolded PI-B Pepper and Gm0970 Spinach2.** Titration of c-di-GMP and cGAMP with tRNA scaffolded PI-B Pepper and Gm0970 Spinach2 biosensors with their respective dye partners (HBC514-Pepper, DFHBI-1T-Spinach2) at varying temperatures (28 ºC –empty circles; 37 ºC – filled circles) with 3 mM Mg2+. (A) Represents the binding affinities of tRNA PI-B Pepper with c-di-GMP or cGAMP, and (B) are the binding affinities of tRNA Gm0970 Spinach2 (Sp2) with c-di-GMP or cGAMP. Points and error bars represent averages and standard deviation of three independent replicates. Best-fit curves are also shown (28 ºC – dotted lines; 37 ºC – solid lines). All binding affinity measurements included 200 nM HBC514 or 10 μM DFHBI-1T with 20 nM RNA. A table of KDs values in nM is displayed in C, where the cognate ligand for each biosensor is bolded.

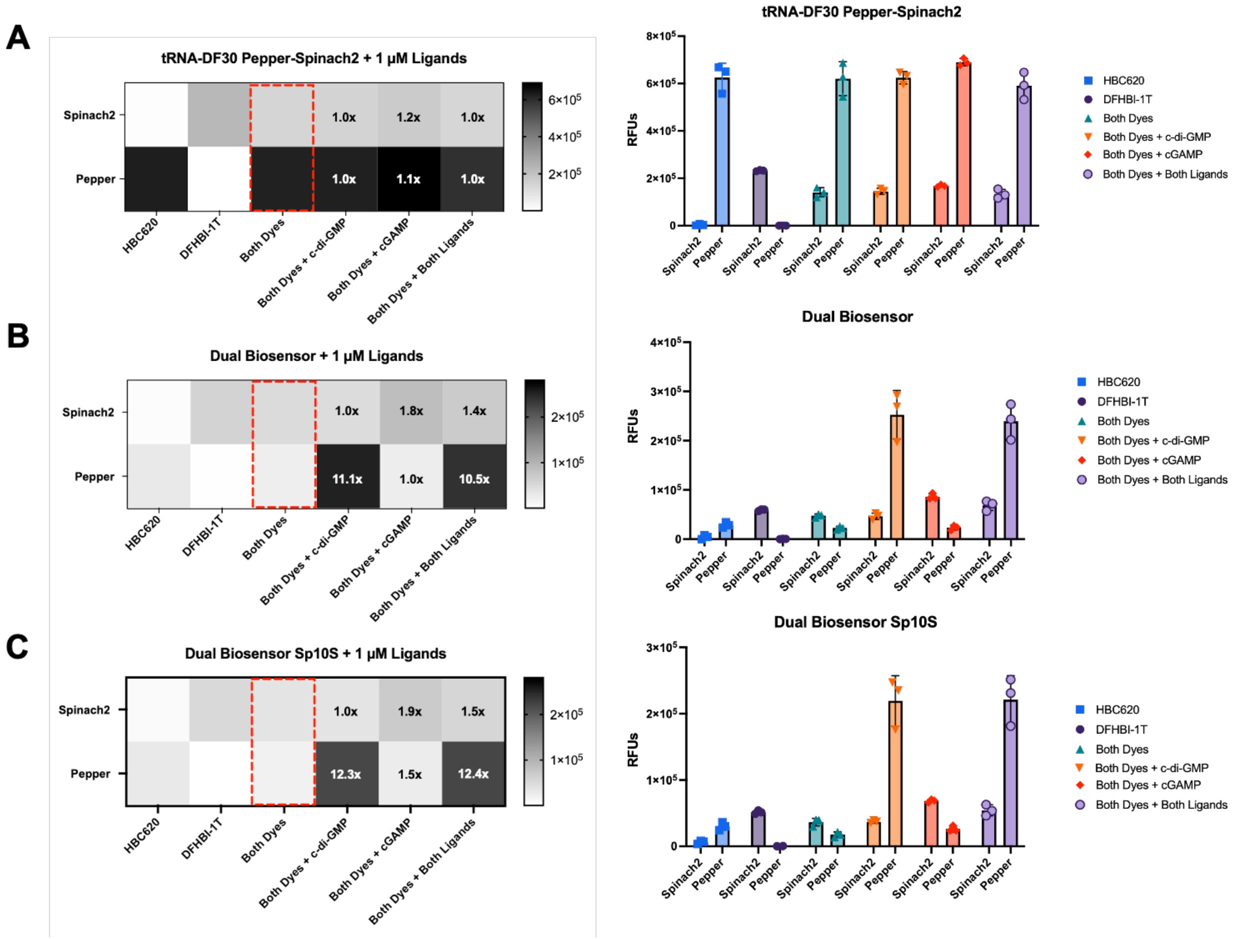


**Supplementary Figure 8. Full *In Vitro* Analysis of Dual Biosensor Designs.** (A-C) Fluorescent activation of tRNA-DF30 Pepper-Spinach2 (control) and biosensor constructs with 1 μM of c-di-GMP and 3'3' cGAMP, along with 10 μM DFHBI-1T and 200 nM HBC620. The left panel represents the results as a heat map, and the right panel shows the raw RFU values in each channel for Pepper (red) and Spinach2 (green) fluorescence under the conditions tested. Fold activation (shown in the left panel) is determined by looking at the fluorescent intensity of Spinach2 and Pepper aptamers with only DFBHI-1T and HBC620 present (red dashed box). All the data shown are the averages of three independent replicates.

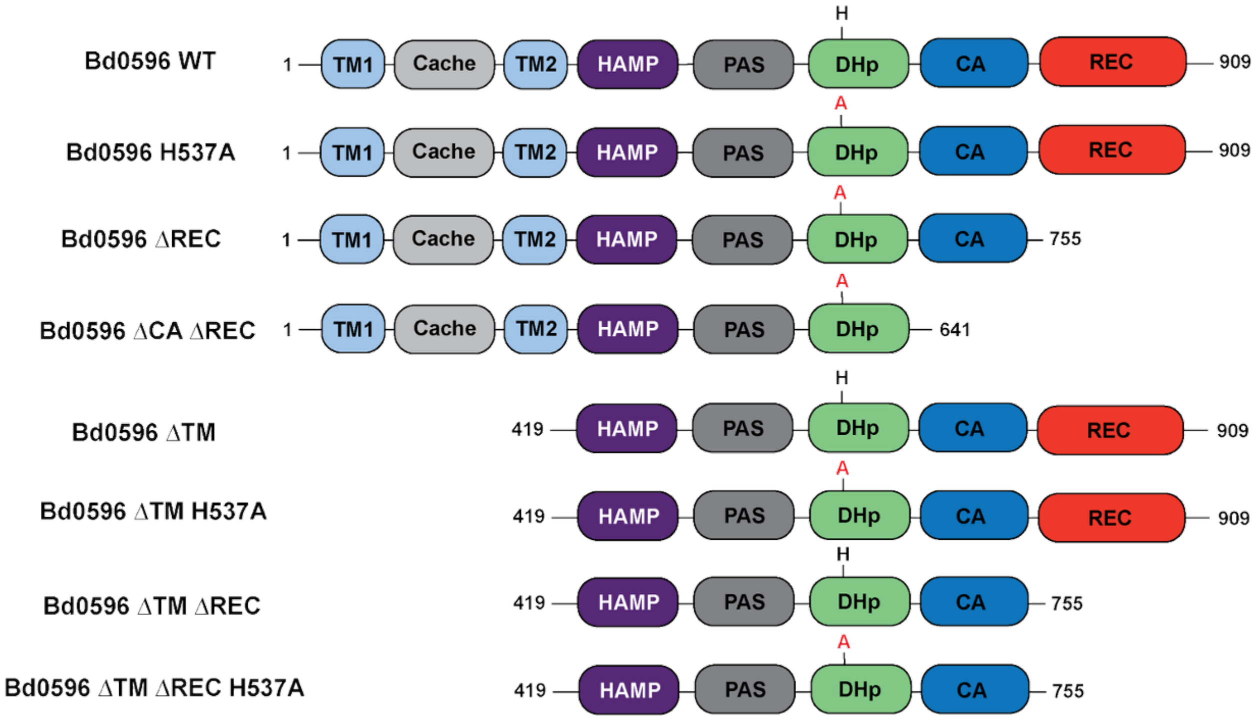


**Supplementary Figure 9. Domain Organization of Bd0596 and Mutant Constructs Used in this Study.** Schematic representation of different Bd0596 constructs used in this study with their boundaries. Position of punctual mutations within different constructs is indicated. TM1 and TM2 stand for transmembrane, DHp for the dimerization and histidine phosphotransfer domain, CA for the ATP binding domain, and REC for the receiver domain.

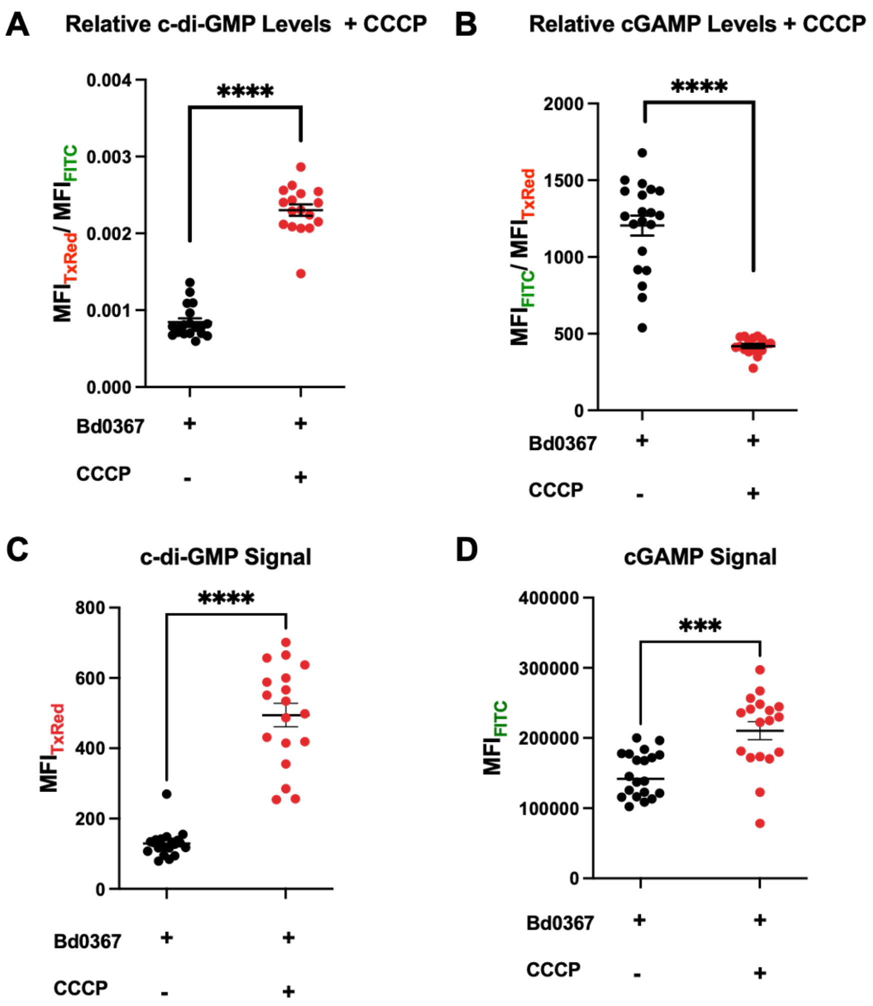


**Supplementary Figure 10. Bd0367 Cyclic Dinucleotide Production is Affected by ATPase Inhibitor, CCCP.** (A-B) Median fluorescence intensities (MFI) are measured via flow cytometry with E. coli BL21 (DE3) Star cells co-expressing Dual Biosensor Sp10S with Bd0367 with or without ATPase inhibitor, CCCP. MFI values of FITC and TxRed channels is normalized to each other to observe the relative c-di-GMP and cGAMP levels within cells. In the left panel are the relative c-di-GMP levels, and in the right panel are the relative cGAMP levels. Data are from 17, 19, or 20 biological replicates by analyzing 50,000 cells per sample after 1 h incubation with DFHBI-1T (50 µM) and HBC620 dye (200 nM). The same biological samples were tested with 5 µM CCCP for 30 min following the 1 h dye incubation. (C-D) Raw MFI values in the TxRed Channel (C) and FITC Channel (D) of Dual Biosensor Sp10S with Bd0367 with or without ATPase inhibitor, CCCP. Each point is the mean of 17, 19, or 20 biological replicates tested by analyzing 50,000 cells per sample. Outliers were identified and excluded from the data using the ROUT method; Q=5 %. Statistical comparison was performed by a two-tailed *t*-test where n=17, 19, or 20. ****P* < 0.001/ *****P* < 0.0001. Data represent the mean ± SEM.

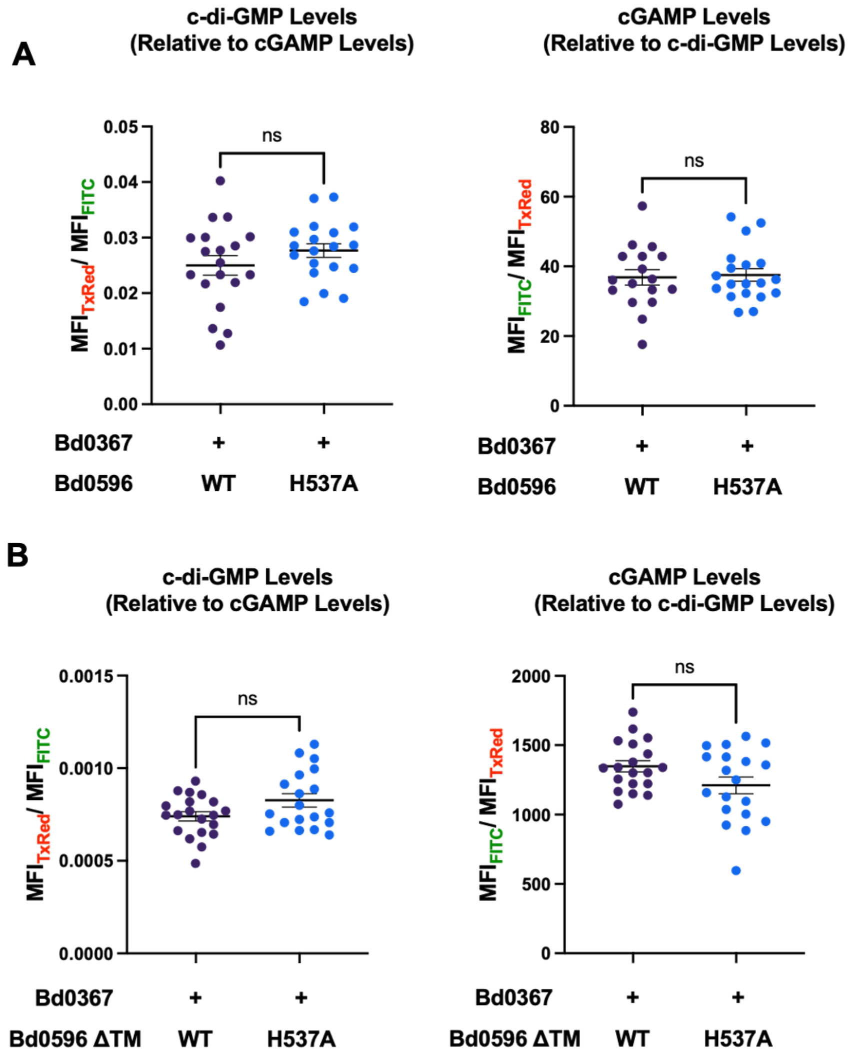


**Supplementary Figure 11. Enzymatic Activity Analysis of Bd0367 with Full-Length and Cytosolic Bd0596.** Median fluorescence intensities (MFI) are measured via flow cytometry with *E. coli* BL21 (DE3) Star cells co-expressing Dual Biosensor Sp10S with various Bd0596 histidine kinase mutants. MFI values of FITC and TxRed channels are normalized to each other to observe the relative c-di-GMP and cGAMP levels within cells. (A) Analysis of full-length Bd0596 and Bd0596 H537A. In the top left panel are the relative c-di-GMP levels, and in the top right panel are the relative cGAMP levels. (B) Analysis of cytosolic Bd0596 ΔTM and Bd0596 ΔTM H537A. In the bottom left panel are the relative c-di-GMP levels, and in the bottom right panel are the relative cGAMP levels. Data are from 19 or 20 biological replicates by analyzing 50,000 cells per sample after 1 h incubation with DFHBI-1T (50 µM) and HBC620 dye (200 nM). Outliers were identified and excluded from the data using the ROUT method; Q=5 %. Statistical comparison was performed by a two-tailed *t*-test where n=19 or 20. ns $P > 0.05$. Data represent the mean ± SEM.

**Supplementary Table 1. PI-B Pepper Biosensor and Dual Metabolite Biosensor DNA Sequences.** The following oligos are DNA template sequences for *in vitro* transcription and to clone into the pET31b plasmid within a tRNA scaffold. In **black** is the tRNA scaffold, underlined is the F30 scaffold, **blue** is the Pepper aptamer sequence, **bold underlined** is the transducer domain sequence, *italic lower case* is the PI-B and Gm0970 riboswitch sequences, **green** is the Spinach2 aptamer sequence, and **purple** is the 10 base-pair spacer sequence. All sequences read from 5' to 3'.

| | |
|---|---|
| PI-B Pepper **1** | CCAATCGTGGCGTGTCGG**A**GATAACGGCAAACTTGTCGAAAGATAAGGACGCAAAGCCACAGGGCCTTCTT<br>GATGAACCGTCAATGGCAGCCTGGCTACC**T**CACTGGCGCCG |
| PI-B Pepper **1.1** | CCAATCGTGGCGTGTCGG**G**GATAACGGCAAACTTGTCGAAAGATAAGGACGCAAAGCCACAGGGCCTTCTT<br>GATGAACCGTCAATGGCAGCCTGGCTACC**C**CACTGGCGCCG |
| PI-B Pepper **2** | CCAATCGTGGCGTGTCGG**AC**GATAACGGCAAACTTGTCGAAAGATAAGGACGCAAAGCCACAGGGCCTTCT<br>TGATGAACCGTCAATGGCAGCCTGGCTACC**GT**CACTGGCGCCG |
| PI-B Pepper **2.1** | CCAATCGTGGCGTGTCGG**AT**GATAACGGCAAACTTGTCGAAAGATAAGGACGCAAAGCCACAGGGCCTTCT<br>TGATGAACCGTCAATGGCAGCCTGGCTACC**AT**CACTGGCGCCG |
| PI-B Pepper **3** | CCAATCGTGGCGTGTCGG**AGC**GATAACGGCAAACTTGTCGAAAGATAAGGACGCAAAGCCACAGGGCCTTC<br>TTGATGAACCGTCAATGGCAGCCTGGCTACC**GCT**CACTGGCGCCG |
| PI-B Pepper **4** | CCAATCGTGGCGTGTCGG**CTT**CGATAACGGCAAACTTGTCGAAAGATAAGGACGCAAAGCCACAGGGCCTT<br>CTTGATGAACCGTCAATGGCAGCCTGGCTACCG**AAG**CACTGGCGCCG |
| Dual Biosensor | GCCCGGATAGCTCAGTCGGTAGAGCAGCGGCCGGTTGCCATGTGTATGTGGGCCAATCGTGGCGTGTCGG<br>*ggataacggcaaacttgtcgaaagataaggacgcaaagccacagggccttcttgatgaaccgtcaatggcagcctggctaccc*CACTGGCGCCGC<br>CCACATACTCTGATGATCCGATGTAACtGAATGAAATGGTGAAGGACGGGTCCAatcgacaatactaaaccatccgcgag<br>ggtgggacggaaagcctacagggtctctgagacagccgggatgccgaaatTTGTTGAGTAGAGTGTGAGCTCCGTAACTAGTTAC<br>ATCGGATCATTCATGGCAACGGCCGCGGGTCCAGGGTTCAAGTCCCTGTTCGGGCGCCA |

| | |
|---|---|
| Dual Biosensor + 10 bp spacer on Sp2 arm | GCCCGGATAGCTCAGTCGGTAGAGCAGCGGCCGGTTGCCATGTGTATGTGGGCCAATCGTGGCGTGTCGG<br>*ggataacggcaaacttgtcgaaagataaggacgcaaagccacagggccttcttgatgaaccgtcaatggcagcctggctaccc*CACTGGCGCCGC<br>CCACATACTCTGATGATCCTGTTGCCATGGATGTAACtGAATGAAATGGTGAAGGACGGGTCCAatcgacaatact<br>aaaccatccgcgagggtgggacggaaagcctacagggtctctctgagacagccgggatgccgaaatTTGTTGAGTAGAGTGTGAGCTCCG<br>TAACTAGTTACATCCATGGCAACAGGATCATTCATGGCAACGGCCGCGGGTCCAGGGTTCAAGTCCCTGTTC<br>GGGCGCCA |

**Table 2. List of Histidine Kinases and Fitness Scores from *B. bacteriovorus* Tn-Seq Screen.** The table is adapted from the full Tn-seq results screen by Duncan et al. Fitness scores (*W*) and standard deviation are from three biological replicates. Fitness values below 0.5 are highlighted in blue, and those above 2.0 are highlighted in orange. $R^2$ values from Bd0367 showcase how closely correlated the histidine kinase knockout strain activity is to the Bd0367 knockout strain. Hypr GGDEF enzyme Bd0367 is highlighted in yellow. Histidine Kinases that are chosen as initial test candidates are highlighted in green.

| H100 Locus | Locus (109J) | ProteinID | VCPL Avg *W* | VCPL Stdv | VCBF Avg *W* | VCBF Stdv | ECPL Avg *W* | ECPL Stdv | ECBF Avg *W* | ECBF Stdv | $R^2$ from Bd0367 |
|---|---|---|---|---|---|---|---|---|---|---|---|
| Bd0367 | EP01_RS13530 | diguanylate cyclase response regulator | 2.2589 | 0.3051 | 1.4397 | 0.2118 | 1.6887 | 0.3159 | 1.6398 | 0.3178 | 0.0000 |
| Bd0596 | EP01_RS16990 | hybrid sensor histidine kinase/response regulator | 2.0122 | 0.2112 | 1.4890 | 0.1585 | 1.4268 | 0.2594 | 1.4690 | 0.2304 | 0.1611 |
| Bd0499 | EP01_RS14130 | hybrid sensor histidine kinase/response regulator | 1.9614 | 0.1818 | 1.6930 | 0.2780 | 1.6546 | 0.1072 | 1.7494 | 0.4656 | 0.165835005 |
| Bd3750 | EP01_RS11025 | two-component sensor histidine kinase | 2.0681 | 0.2500 | 1.6150 | 0.0716 | 2.0737 | 0.1095 | 1.6175 | 0.1380 | 0.215884292 |
| Bd0584 | EP01_RS16935 | hybrid sensor histidine kinase/response regulator | 1.8210 | 0.6623 | 1.6490 | 0.3641 | 1.6382 | 0.4906 | 1.4592 | 0.1807 | 0.270777343 |
| Bd0300 | EP01_RS13220 | hybrid sensor histidine kinase/response regulator | 1.8227 | 0.0719 | 1.4529 | 0.1192 | 1.4429 | 0.0574 | 1.4736 | 0.1068 | 0.278503794 |
| Bd2184 | EP01_RS07085 | signal transduction sensor histidine kinase | 1.8424 | 0.1540 | 1.7087 | 0.3640 | 1.9333 | 0.4509 | 1.6634 | 0.6286 | 0.306279874 |
| Bd2843 | EP01_RS10160 | hybrid sensor histidine kinase/response regulator | 1.9471 | 0.0972 | 1.8775 | 0.2963 | 1.5394 | 0.1314 | 1.6237 | 0.3131 | 0.311405843 |
| Bd1382 | EP01_RS16535 | hybrid sensor histidine kinase/response regulator | 2.1116 | 0.3196 | 1.9543 | 0.3854 | 1.7824 | 0.2204 | 1.9086 | 0.2074 | 0.3676 |
| Bd1018 | EP01_RS14910 | PAS domain-containing sensor histidine kinase | 1.7239 | 0.1565 | 1.4255 | 0.2774 | 1.9544 | 0.3927 | 1.4878 | 0.0882 | 0.380166243 |
| Bd3779 | EP01_RS11195 | two-component sensor histidine kinase | 1.6473 | 0.1342 | 1.4381 | 0.1150 | 1.6092 | 0.1408 | 1.5610 | 0.1661 | 0.38665232 |
| Bd1758 | EP01_RS04815 | two-component sensor histidine kinase | 1.7005 | 0.5356 | 1.4723 | 0.8989 | 1.5471 | 0.3617 | 1.3301 | 0.2745 | 0.428909789 |
| Bd3367 | EP01_RS02005 | ATP-binding protein | 1.6374 | 0.3817 | 1.4371 | 0.3382 | 1.4278 | 0.5653 | 1.5641 | 0.2620 | 0.460071229 |

| | | | | | | | | | | | |
|---|---|---|---|---|---|---|---|---|---|---|---|
| Bd2849 | EP01_RS10190 | histidine kinase | 1.6056 | 0.2404 | 1.4045 | 0.2182 | 1.6433 | 0.2795 | 1.2778 | 0.1948 | 0.561221306 |
| Bd3528 | EP01_RS02775 | histidine kinase | 1.5750 | 0.3260 | 1.5001 | 0.1464 | 1.3648 | 0.3113 | 1.5149 | 0.2897 | 0.591934414 |
| Bd2142 | EP01_RS06890 | hypothetical protein | 1.6317 | 0.1169 | 1.7736 | 0.2933 | 2.0058 | 0.2326 | 1.7121 | 0.4167 | 0.610721292 |
| Bd3305 | EP01_RS01735 | hybrid sensor histidine kinase/response regulator | 1.4852 | 0.4143 | 1.4232 | 0.7895 | 1.6202 | 0.5383 | 1.4612 | 0.4786 | 0.635491717 |
| Bd0578 | EP01_RS16905 | histidine kinase | 2.1182 | 0.2162 | 2.0756 | 0.2138 | 2.0203 | 0.2339 | 1.9935 | 0.1227 | 0.659274732 |
| Bd3527 | EP01_RS02770 | response regulator | 2.0885 | 0.3179 | 2.0553 | 0.7070 | 2.1241 | 1.0232 | 1.9027 | 0.3572 | 0.666770383 |
| Bd3430 | EP01_RS02300 | histidine kinase | 2.5317 | 0.3848 | 2.0745 | 0.1988 | 2.0456 | 0.0999 | 1.9687 | 0.2375 | 0.712900588 |
| Bd3613 | EP01_RS03150 | histidine kinase | 1.4464 | 0.2056 | 1.1971 | 0.1135 | 1.6134 | 0.1163 | 1.3003 | 0.2647 | 0.840024858 |
| Bd3648 | EP01_RS10625 | PAS domain-containing sensor histidine kinase | 1.9467 | 0.3251 | 2.0579 | 0.1566 | 2.1183 | 0.4085 | 2.0613 | 0.4726 | 0.841936053 |
| Bd2406 | EP01_RS08135 | histidine kinase | 2.5055 | 0.0885 | 2.1618 | 0.4996 | 2.3243 | 0.2299 | 1.8832 | 0.1785 | 1.045465472 |
| Bd3359 | EP01_RS01975 | histidine kinase | 2.2884 | 1.0289 | 2.1502 | 0.4758 | 2.4516 | 0.3318 | 2.0549 | 0.2600 | 1.260040341 |
| Bd1828 | EP01_RS05140 | hybrid sensor histidine kinase/response regulator | 1.8213 | 0.6069 | 2.3950 | 0.8158 | 2.0483 | 0.7764 | 1.9122 | 0.5899 | 1.3077 |
| Bd2576 | EP01_RS08865 | two-component sensor histidine kinase | 1.4708 | 0.6905 | 1.1637 | 0.5435 | 1.0389 | 0.4630 | 1.2031 | 0.5796 | 1.310185222 |
| Bd2584 | EP01_RS08900 | PAS domain-containing sensor histidine kinase | 1.2337 | 0.1614 | 1.2910 | 0.0361 | 1.2704 | 0.1407 | 1.3660 | 0.2545 | 1.323143539 |
| Bd2833 | EP01_RS10115 | histidine kinase | 1.2470 | 0.0838 | 1.3873 | 0.2773 | 1.1830 | 0.0597 | 1.2282 | 0.1837 | 1.451890419 |
| Bd1335 | EP01_RS16295 | histidine kinase | 1.1483 | 0.2685 | 1.3564 | 0.3555 | 1.2457 | 0.0719 | 1.3215 | 0.2065 | 1.537877503 |
| Bd1657 | EP01_RS04390 | PAS domain-containing sensor histidine kinase | 1.0143 | 0.3242 | 1.4813 | 0.3445 | 0.9088 | 0.0254 | 1.4721 | 0.1690 | 2.18716155 |

| | | | | | | | | | | | |
|---|---|---|---|---|---|---|---|---|---|---|---|
| Bd0216 | EP01_RS12815 | two-component sensor histidine kinase | 0.9102 | 0.0446 | 1.0121 | 0.1845 | 0.7154 | 0.1380 | 1.1269 | 0.2172 | 3.212105242 |
| Bd1855 | EP01_RS03820 | hybrid sensor histidine kinase/response regulator | 0.9734 | 0.2107 | 0.6710 | 0.1766 | 1.0917 | 0.2195 | 0.8436 | 0.2976 | 3.233790364 |
| Bd1506 | EP01_RS03680 | histidine kinase | 0.6003 | 0.2454 | 0.4912 | 0.5351 | 0.3920 | 0.1373 | 0.7017 | 0.2186 | 6.212110171 |
| Bd3393 | EP01_RS02120 | two-component sensor histidine kinase | 0.4489 | 0.1039 | 0.6189 | 0.1068 | 0.4556 | 0.1151 | 0.5556 | 0.0875 | 6.645801273 |
| Bd3125 | EP01_RS00910 | GGDEF domain-containing protein | 0.3149 | 0.1058 | 0.5561 | 0.1722 | 0.4153 | 0.0623 | 0.6913 | 0.0728 | 7.080996901 |
| Bd3126 | EP01_RS00915 | two-component sensor histidine kinase | 0.0723 | 0.0627 | 0.6969 | 0.7827 | 0.5220 | 0.2250 | 0.4397 | 0.4237 | 8.134293987 |
| Bd3428 | EP01_RS02295 | histidine kinase | 0.0899 | 0.0780 | 0.0382 | 0.0333 | 0.0000 | 0.0000 | 0.1099 | 0.0956 | 11.86124817 |
| Bd0620 | EP01_RS17090 | two-component sensor histidine kinase | 0.0000 | 0.0000 | 0.0545 | 0.0944 | 0.0286 | 0.0495 | 0.0913 | 0.1500 | 12.17524297 |
| Bd1512 | EP01_RS03710 | pilS sensor protein | 0.0000 | 0.0000 | 0.0000 | 0.0000 | 0.0000 | 0.0000 | 0.0155 | 0.0268 | 12.66549466 |
| Bd1899 | EP01_RS18290 | two-component sensor histidine kinase | 6.6537 | 4.7175 | 6.1002 | 4.0065 | 5.1931 | 3.4775 | 5.1987 | 3.7376 | 65.9811671 |
| Bd3036 | EP01_RS00500 | PAS domain-containing sensor histidine kinase | ND | ND | ND | ND | ND | ND | ND | ND | N/A |
| Bd3214 | EP01_RS01335 | hypothetical protein | ND | ND | ND | ND | ND | ND | ND | ND | N/A |
| Bd3450 | EP01_RS02385 | two-component sensor histidine kinase | ND | ND | ND | ND | ND | ND | ND | ND | N/A |
